# Resting natural killer cells promote the progress of colon cancer liver metastasis by elevating tumor-derived sSCF

**DOI:** 10.1101/2024.02.27.582307

**Authors:** Chenchen Mao, Yanyu Chen, Dong Xing, Teming Zhang, Yangxuan Lin, Cong Long, Jiaye Yu, Yunhui Luo, Tao Ming, Wangkai Xie, Zheng Han, Dianfeng Mei, Dan Xiang, Mingdong Lu, Xian Shen, Xiangyang Xue

## Abstract

**Purpose:** The abundance and biological contribution of Natural killer (NK) cells in cancer are controversial. Here, we aim to uncover clinical relevance and cellular roles of NK cells in colon cancer liver metastasis (CCLM)

**Methods:** We integrated single-cell RNA sequencing, spatial transcriptomics, and bulk RNA-sequencing datasets to investigate NK cells’ biological properties and functions in the microenvironment of primary and liver metastatic tumors. Results were validated through an in vitro co-culture experiment based on bioinformatics analysis.

**Results:** We used single-cell RNA sequencing and spatial transcriptomics to map the immune cellular landscape of colon cancer and well-matched liver metastatic cancer. We discovered that GZMK+ resting NK cells increased significantly in tumor tissues and were enriched in the tumor regions of both diseases. After combining bulk RNA and clinical data, we observed that these NK cell subsets contributed to a worse prognosis. Meanwhile, KIR2DL4+ activated NK cells exhibited the opposite position and relevance. Pseudotime cell trajectory analysis revealed the evolution of activated to resting NK cells. In vitro experiments further confirmed that tumor-cell-co-cultured NK cells exhibited a decidual-like status, as evidenced by remarkable increasing CD9 expression. Functional experiments finally revealed that NK cells exhibited tumor-activating characteristics by promoting the dissociation of SCF (stem cell factor) on the tumor cells membrane depending on cell-to-cell interaction, as the supernatant of the co-culture system enhanced tumor progression.

**Conclusion:** Together, our findings revealed a population of protumorigenic NK cells that may be exploited for novel therapeutic strategies to improve therapeutic outcomes for patients with CCLM.

## Introduction

Colon cancer is one of the most common malignancies, with more than 1 million new cases and 500 thousand deaths reported globally in 2020(1). Approximately 19– 26% of patients with colon cancer present with synchronous metastatic diseases at the first diagnosis(2), among which 14.5–17.5% develop liver metastasis(2, 3). Although progress in the treatment of metastatic disease, including improved diagnosis and treatment strategies for liver metastases(4–6), increased cancer-directed surgery(7), and the development of targeted therapies(8), has greatly improved the survival of these patients in recent decades, the 2-year relative survival rate for patients diagnosed with distant-stage disease was 36%(2), significantly lower than those without metastasis. Thus, colon cancer-derived liver metastasis (CCLM) is a clinical challenge that requires urgent development of novel treatment methods.

The tumor microenvironment (TME) is a dynamic environment that governs tumor behavior and is required for tumor cell survival, growth, proliferation, and metastasis(9). Unlike previous understanding, it has recently been proposed that certain immune cells present in the TME (such as myeloid-derived suppressor cells(10), macrophages(11), neutrophils(12), and CD8^+^ T cells(13)) favor tumor progression. Thus, the contradictory findings of the pro- and anti-tumor effects of tumor-infiltrating immune cells require a better understanding of the immune features and profile heterogeneity of colon cancer liver metastasis (CCLM) for the future development of immune-modulatory strategies to stratify and target immune cells.

Natural killer (NK) cells, an important component of tumor-infiltrating immune cells, are cytotoxic lymphocytes that play a key role in recognizing and eliminating malignant cells and are therefore considered the early responders against tumors(14). Among various immune cells, tumor-infiltrating NK cells are associated with prognosis in various cancers, such as colorectal cancer (CRC)(15), lung cancer(16), and gastric cancer(17). Recently, based on their anti-tumor potential, NK cells were exploited for treating malignancies and have proven to be highly promising for the treatment of certain hematologic malignancies(18, 19) but have limited efficacy for treating solid tumors(20–22). Additionally, intratumor NK cells differ phenotypically or functionally from peripheral NK cells in several malignancies(23, 24); a specific subset of tumor-infiltrating NK cells (CD11b^−^ and CD27^−^) as well as decidual-like NK (CD9^+^ and cd49a^+^) was associated with tumor progression in lung and hepatocellular cancers(25–27). Notably, the tumor-promoting transformation of NK cells may be educated by cancer cells(28, 29). However, their function remains elusive. In particular, the presence of specific NK cell subsets within colon cancer and their potential relationship to CCLM progression have not been fully characterized to date.

In this study, we combined previously published single-cell RNA sequencing, spatial transcriptomics (ST), and bulk RNA-sequencing data from public datasets, aiming to comprehensively chart the cellular landscape of TME in primary colon cancer and matched liver metastasis. Results of bioinformatics analysis were further validated through an in vitro co-culture experiment (Fig. 1A).

**Figure 1.**
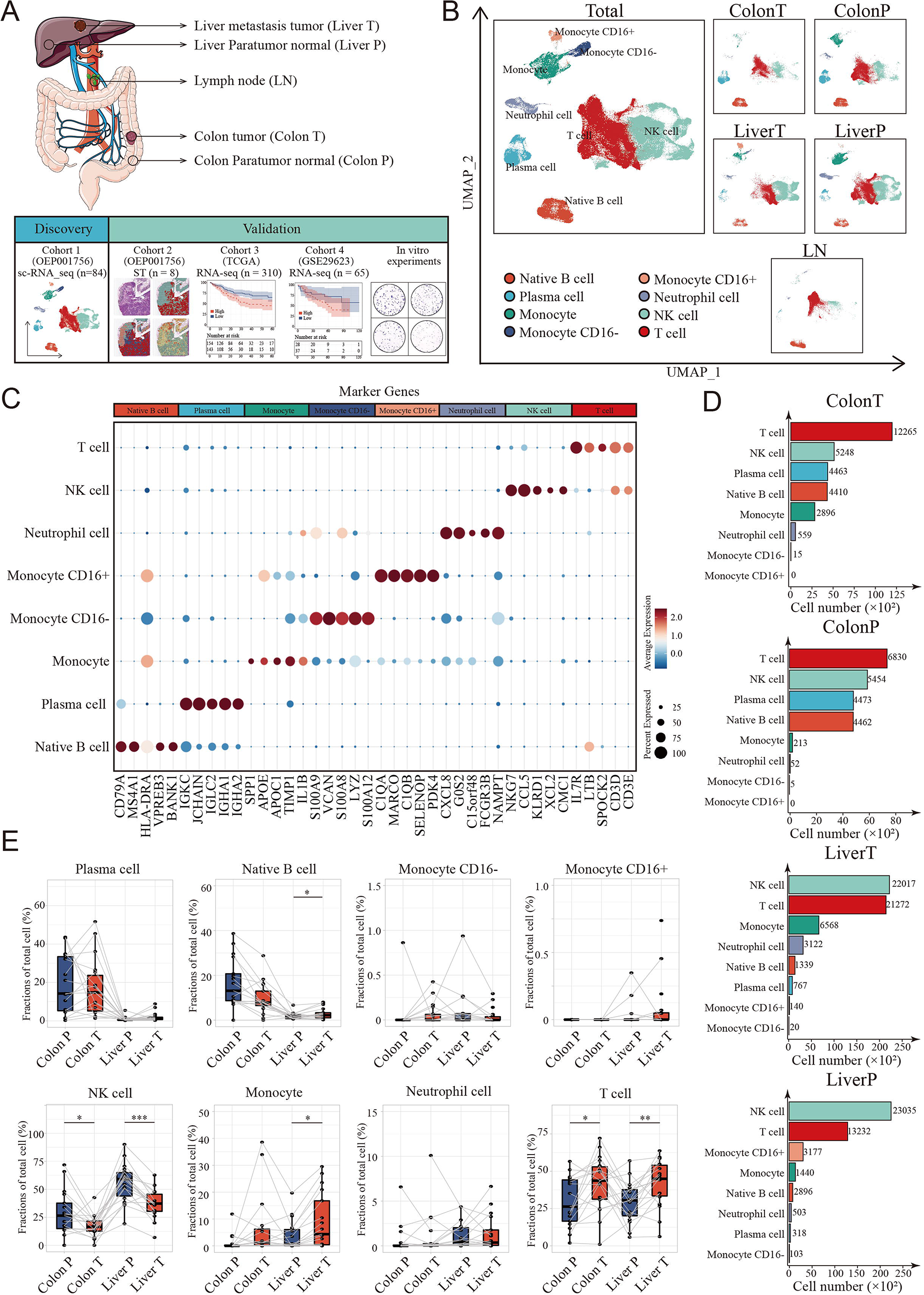
Landscape of tumor immune microenvironment in CCLM revealed by single-cell transcriptomics. A. Schematic overview of the experimental design and analytical workflow. Colon T: colon tumor tissue; Colon P: colon paratumor normal tissue; Liver T: liver metastasis tumor tissue; Liver P: liver paratumor normal tissue; LN: lymph node metastasis tissue. B. The UMAP plot of all main immune cell types. C. Dot plots showing average expression of highly variable genes of each cell group. The dot size represents percent of cells expressing the genes in each cluster and the color of dot represents the expression intensity. D. The cell numbers of main immune cells across tissues. E. Proportions of all main immune cells. P values were determined by the paired nonparameter test.

## Materials and Methods

### Data sources

In this study, we integrated four independent public datasets that contained single-cell RNA sequencing and ST data (10X Genomics) (derived from previously published research at the website: http://www.cancerdiversity.asia/scCCLM/)(30). And two bulk transcriptomics data of colon cancer (downloaded from TCGA cohort COAD at the website: https://xena.ucsc.edu/ and NCBI’s Gene Expression Omnibus with accession number GSE29623).

### Single cell RNA Sequencing Analysis

The R (v4.0.5) package Seurat (v4.1.0)(31) was used to process the scRNA data. Since dataset quality control had been performed and Seurat objects had been created in previous studies, we did not further filter the scRNA-seq data or remove the batch effects. The SCTransform method was used to normalize the data. After selecting 2,000 highly variable genes using the Find Variable Features function in Seurat, principal component analysis was performed using these genes to reduce the data dimensions. Based on the ElbowPlot function in Seurat, we chose the top 30 principal components to run the FindNeighbors function. Next, the cells were clustered using the FindClusters function with a resolution of 0.1 for clustering immune cells. A uniform manifold approximation and projection algorithm was used for data visualization, as previously reported(32). Differentially expressed genes (DEGs) of each subset were identified using the FindAllMarkers function in Seurat. SingleR (v1.0.0) was used to name each cluster(33). Additionally, the subgroup of NK cells was defined by the preferential expression marker of the resting and activated NK cells described in the CIBERSORT.

### Pseudotime Analysis

To analyze the differentiations of NK cells, monocle2 (http://cole-trapnell-lab.github.io/monocle-release), which uses an algorithm to learn the changes in gene expression sequences that each cell must undergo as part of a dynamic biological process(34), was used for pseudotime trajectory analysis to identify the transitional relationships among different clusters. The cells were reduced dimensionally using the DDRTree method, sequenced in pseudotime, and finally visualized(35).

### Bulk RNA seq Data Analysis

The bulk transcriptome RNA-seq data and corresponding clinical data were obtained from The Cancer Genome Atlas of colon adenocarcinoma through the UCSC Xena browser (GDC hub) (https://gdc.xenahubs.net)(36). In total, 459 colon cancer samples with survival information were downloaded, and 310 samples were finally enrolled (11 formalin-fixed samples were excluded, and an additional 134 and 4 samples were excluded due to deletion of metastasis information and sequencing matrix, respectively). Transcriptomic data from 65 colon cancer samples in GSE29623 were obtained as the validation cohorts. CIBERSORT(37), xCELL(38), EPIC(39), and MCP-counter(40), which use gene expression to infer the proportions of tumor-infiltrating immune cells, were used to analyze the TME cell type. R packages survival (v3.2-10) and survminer (v0.4.9) were used for survival analysis. The Youden index was selected as the cutoff value to differentiate patients into distinct groups (high or low). The Kaplan–Meier survival curve was modeled using the survfit function. Subsequently, a Cox proportional hazards regression model was established to determine the independent risk factors. DEGs between the metastasis and non-metastasis groups were determined with the filtering condition of log2 |fold change|>1 and p-value <0.05, using the R package limma. GO and KEGG pathway functional enrichment analyses were performed using the clusterProfiler R package (v3.18.1) to assign various biological processes, molecular functions, cellular components, and pathways of identified marker genes in the cluster of interest(41), and p <0.05 was regarded as statistically enriched. To explore the different KEGG pathways and hallmark gene sets between the metastasis and non-metastasis groups, gene set enrichment analysis (GSEA) was performed using data from The Molecular Signatures Database (c2.cp.kegg.v7.3.symbols) and the fgsea R package (v1.12.0)(42). Pathways with an adjusted p-value below 0.05 were deemed to be significantly enriched.

### ST Data Analysis

Seurat was also used for ST data processing and visualization. We used the SCT method to normalize the ST data; the functions SelectIntegrationFeatures, PrepSCTIntegration, FindIntegrationAnchors, and IntegrateData were used to integrate the ST data. An unsupervised clustering method was subsequently used to cluster similar ST spots. Cell population annotations were based on hematoxylin and eosin (H&E) staining sections and the highly variable genes in each cluster. Scores of cell-specific signatures (top 20 DEGs) from scRNA-seq were calculated using the AddModuleScore function. SpatialDimPlot and SpatialFeaturePlot were combined to visualize the cell expression level in the ST data(32). Since the tumor and non-tumor region was identified by immunohistochemistry as well as cell clusters in ST data, the differences of cell subgroup presence in tumor and non-tumor region were determined by visual inspection

### Cell Lines and Co-cultures

Colon carcinoma cell lines HCT-116, DLD-1and Human Foreskin Fibroblast (HFF) were purchased from the Cell Bank of the Chinese Academy of Sciences (Shanghai, China). HCT-116, DLD-1 were grown in a RPMI-1640 medium (GIBCO/BRL) supplemented with 10% FBS (GIBCO/BRL) while HFF were grown in the DMEM+10% FBS. The human NK cell line NK-92 was also purchased from the Cell Bank of the Chinese Academy of Sciences (Shanghai, China) and maintained in RPMI-1640 supplemented with 12.5% horse serum (GIBCO), 12.5% FBS, 100 U/mL rhIL-2, 0.1 mmol/L b-mercaptoethanol, and 0.02 mmol/L folic acid. All the cells were cultured in a humidified chamber at 37 °C and 5% CO_2_.

Colon cancer cells were plated at a density of 1×10^5^ cells/well in 6-well plates for 24 h. Supernatants were collected and cells and debris therein were removed by centrifugation. NK cells were co-cultured with cancer cells in a ratio of 1:1 in a fresh mixed medium for a further 24h (CN group). Additionally, NK cells were also cultured in a mixed medium of supernatant and NK medium in the ratio 1:1(SN group) as well as fresh RPMI-1640 10% FBS and NK medium in the same ratio (MN group) for 24h (Fig.S1A). When indicated, co-cultures were performed in Transwell devices (JET biofil), maintaining the same ratios and culture times. Finally, as shown in Fig.S1B, the co-cultured supernatant of CN, SN and MN group were collected, which determined as CNS, SNS and MNS, to incubate fresh HCT-116 to explore the NK cell-mediated inductive effect on colon cells.

### Flow Cytometry Analysis

NK cells in each co-cultured group were collected and washed twice in 1 × PBA. Then, they were incubated with APC-labeled CD56 (Biolegend), PE-labeled KIR2DL4 (Affinity Biosciences), FITC-labeled GZMK (Affinity Biosciences), FITC-labeled CD9 (Biolegend), PE-labeled PD-1 (Biolegend) and PerCP-labeled CD49a antibody for 30 min, rewashed, and resuspended in PBS. Fluorescence Minus One (FMO) control was set up to define the position of the positive gate as shown in Table. S1 and Table. S2. BD flow cytometry was used for detection.

### Cell Counting Kit-8 (CCK-8) Cell Viability and Cell Colony Formation Assay

Colon cancer cells (5,000 cells/well) were cultured in 96-well plates with co-cultured supernatants from different co-culture groups alone or with both co-cultured supernatants and imatinib mesylate (2 µM) as previously reported(43) for 24 h. At pre-determined time points, 10 μL of CCK-8 reagent (Dojindo, Japan) was added and incubated for 2 h at 37 °C, and then the absorbance was measured using a microplate reader (Thermo Scientific) at 450 nm. All experiments were carried out in triplicate.

For the cell colony formation assay, colon cancer cells (500 cells/well) were seeded in 12LJwell plates with co-cultured supernatants alone or with both co-cultured supernatants and imatinib mesylate at 37 °C for 1 week. Then, cell colonies were fixed with 4% paraformaldehyde for 10 min and stained with 0.5% crystal violet for 5 min. Cell colonies containing >20 cells were counted. All experiments were carried out in triplicate.

### Transwell Migration/Invasion Assays

The polycarbonate membrane in the transwell chambers was coated with Matrigel (Corning, NY, USA). Next, we transferred 1 × 10^5^ cells from the serum-free medium with or without imatinib mesylate into the top chamber, added co-cultured supernatants alone or with both co-cultured supernatants and imatinib mesylate in the bottom chamber, and incubated at 37 °C for 24 h. Then, we removed the non-invading cells on the top side of the membrane by scrubbing, fixed the migrating or invading cells at the bottom side of the membrane with 4% paraformaldehyde, and stained with 0.5% crystal violet. The number of cells was counted under a microscope (Leica, London, UK) from four randomly chosen fields per well to determine the number of cells in each group.

### Total RNA Extraction and Quantitative Real-Time PCR (qPCR)

Total RNA was extracted from cells using TRIZOL reagent (Invitrogen, USA). RNA was reverse transcribed into cDNA using a PrimeScript RT Reagent Kit (Takara, Japan). qPCR was performed using QuantStudio™ Test Development Software (Thermo Scientific, Waltham, MA, USA) with SYBR Green qPCR Master Mix (EZBioscience, Roseville, MN, USA). The sequence of KITLG and housekeeping gene GAPDH primers is listed in Table S3. KITLG mRNA data were normalized to that of GAPDH.

### Luminex Liquid Suspension Chip Detection and ELISA

Luminex liquid suspension chip detection was performed by Wayen Biotechnologies (Shanghai, China). The Bio-Plex Pro Human Chemokine Panel 48-plex kit was used in accordance with the manufacturer’s instructions. Briefly, supernatants from different co-cultured groups were incubated in 96-well plates embedded with microbeads for 1 h and then incubated with a detection antibody for 30 min. Subsequently, streptavidin-PE was added in each well for 10 min, and values were read using the BioPlex MAGPIX System (Bio-Rad). The Human SCF ELISA kit (Solarbio, China) was used according to the manufacturer’s instructions.

### Statistical Analysis and Visualization

All statistical analyses were performed using SPSS (v23.0; IBM SPSS, Chicago, IL, USA) and R (v4.0.5), and data visualization was performed on R packages Seurat (v4.1.0), ggplot2 (v3.3.5), ggsignif (v0.6.1), and pheatmap (v1.0.12).

## Results

### Integrated scRNA-seq and ST Precisely Quantified Immune Cell Diversity in CCLM

We used a previously published single-cell dataset containing 89 samples from paired samples of colon cancer, adjacent colon, liver metastasis, adjacent liver, lymph nodes along colons, and peripheral blood mononuclear cells from 20 patients to define the heterogeneous immune microenvironment landscapes of CCLM. Subsequently, 178,630 CD45+ cells were integrated.

To further define the main cell type, clustering analysis and SingleR were performed, and monocytes, neutrophil cells, native B cells, plasma cells, T cells, and NK cells were identified from the CCLM samples (Fig. 1B-C; Fig. S2A and B; Table.S4). Notably, T and NK cells were the major components of all immune cells and distinct across different tissue types, with NK cells significantly decreasing in tumor tissues (both primary colon cancer and liver metastasis cancer) compared to corresponding adjacent normal tissues, whereas T cells were the opposite (Fig. 1D and E; Fig. S2C).

To comprehensively analyze the spatial distribution profile of colon primary tumors and CCLM tumors, we collected ST data, which came from eight samples including paired primary colon cancer and liver metastasis. Unsupervised clustering analysis was first performed to cluster similar ST spots, and the annotation of the clusters was further determined according to cell marker genes (Fig. S3A-C, Table.S5). Further, after combining the gene expression features of each sample (Fig. S4) and H&E staining, six morphological regions, including tumor, fibroblast, smooth muscle, B cells, hepatocytes, normal epithelium, and tumor and paratumor areas, were identified. (Fig. 2A-C).

**Figure 2.**
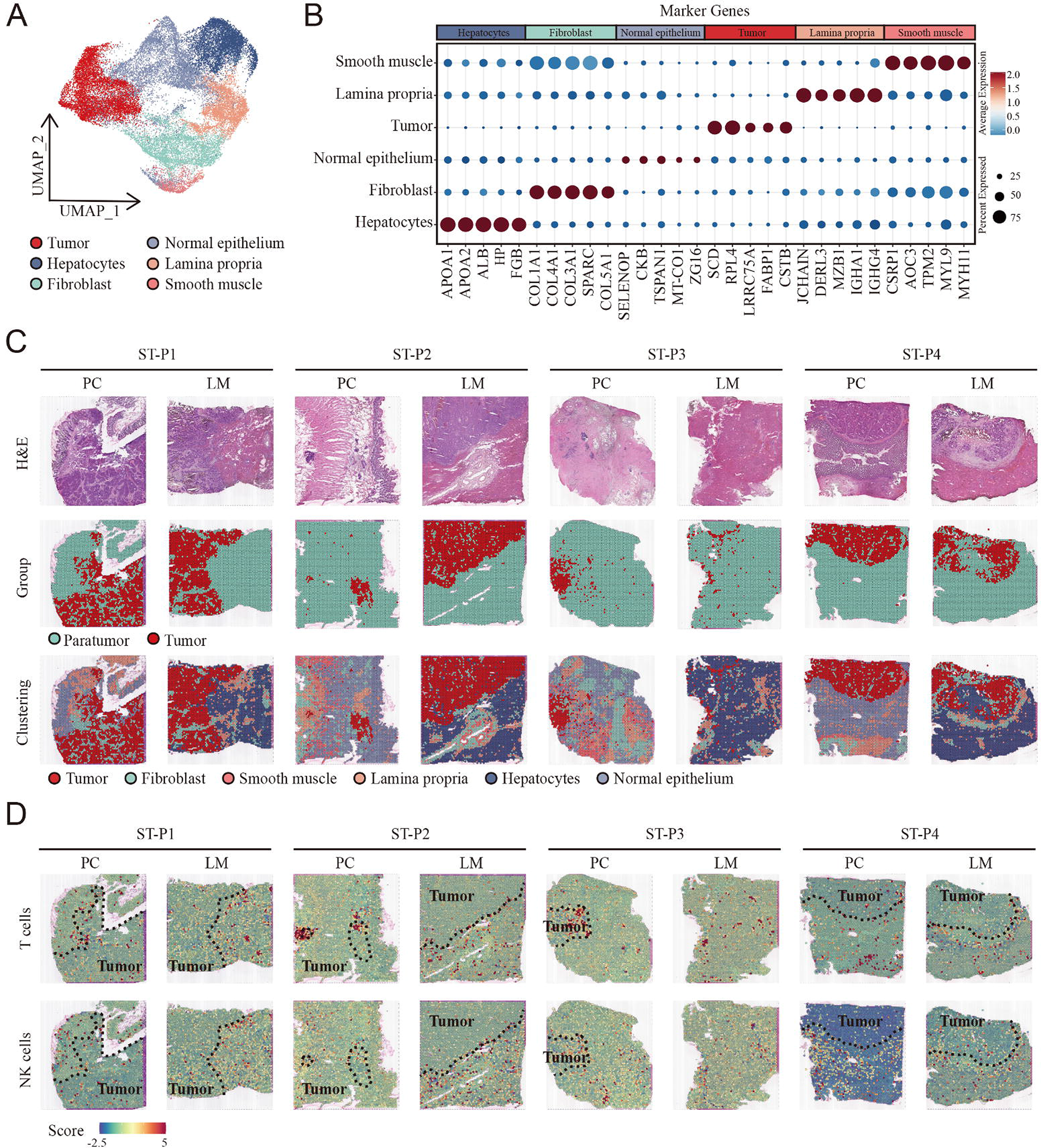
Cellular identification in spatial transcriptomic samples. A. UMAP visualization of cell clusters in spatial transcriptomic samples. B. Dot plots showing average expression of markers in indicated cell clusters. C. Overview of the spatial transcriptomic sections. H&E staining of spatial transcriptomic sections (upper). Tumor tissue and paratumor tissue identification of each section (middle). Spatial cluster distribution of each section (lower). PC: primary cancer; LM: liver metastasis. ST-P1: spatial transcriptomic dataset patient 1, ST-P1: spatial transcriptomic dataset patient 1; ST-P2: spatial transcriptomic dataset patient 2; ST-P3: spatial transcriptomic dataset patient 3; ST-P4: spatial transcriptomic dataset patient 4. D. The signature scores of T cells (upper) and NK cells (lower) in colon cancer and liver metastasis in the spatial transcriptomic sections.

To integrate the scRNA-seq and ST data, we used the AddModuleScore function in Seurat to quantify the main immune subpopulations. Consistent with scRNA-seq, T cells were remarkably enriched in the tumor region of both primary colon cancer and liver metastasis cancer, whereas NK cells were enriched in the non-tumor area (Fig. 2D).

### Biological Relationship Between NK Cells and Metastasis of Colon Cancer Revealed by Bulk RNA Transcriptomics

TME cell composition differences between metastasis and non-metastasis colon cancer were evaluated using multiple tools that could robustly quantify the abundance of cell populations based on transcriptomic datasets, including xCell, EPIC, quanTIseq, and MCPcounter and bulk-seq datasets (TCGA COAD cohort). The percentages of NK cells markedly decreased in metastasis colon cancer using MCPcounter, EPIC, and xCell (Fig. 3A-D).

**Figure 3.**
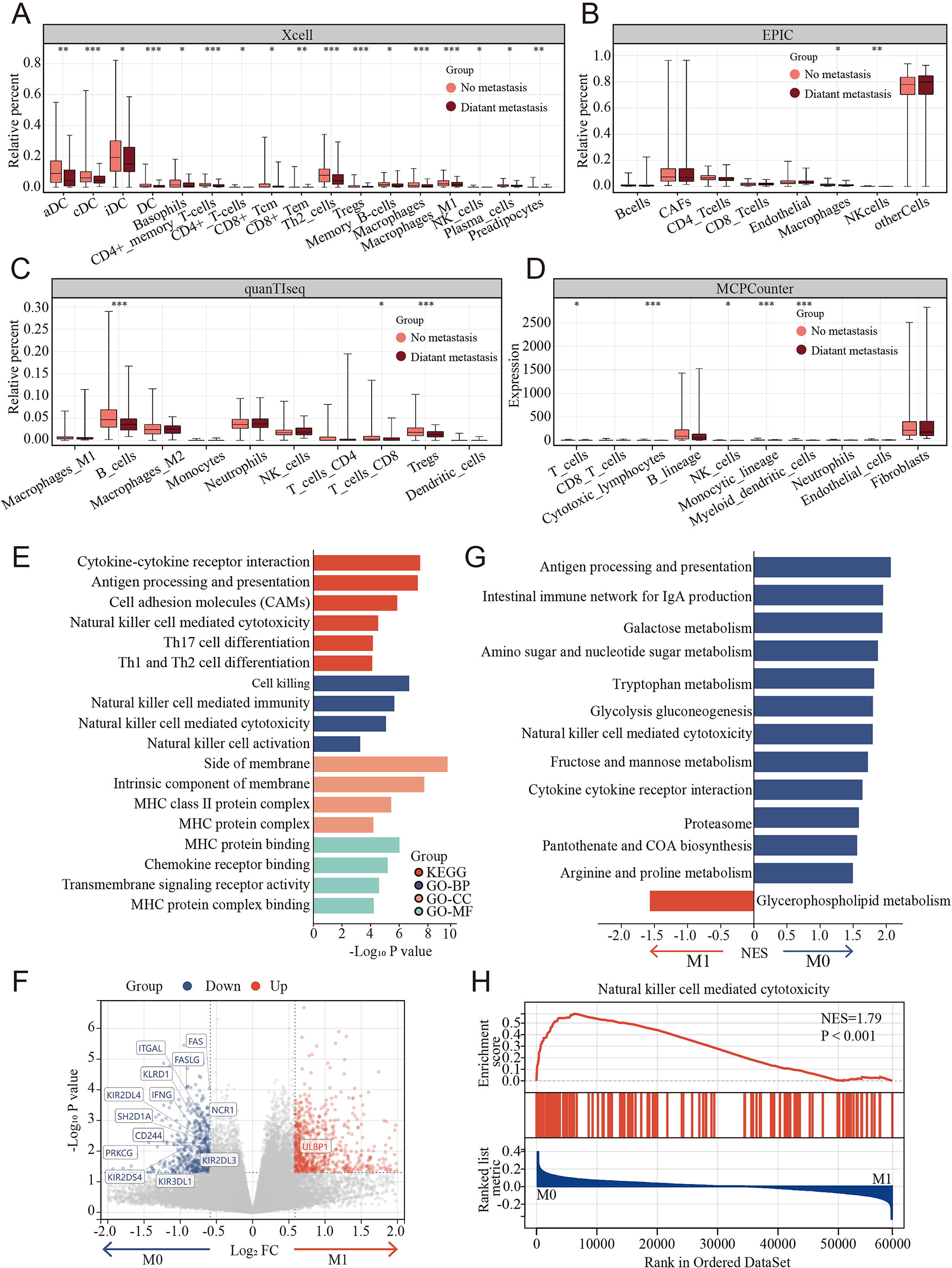
Clinical and biological relationship between NK cells and metastasis of colon cancer revealed by Bulk RNA transcriptomics. A-D. The relationship of immune cell percentage determined by xCell, EPIC, quanTIseq and MCPCounter between metastasis and non-metastasis tumor in TCGA COAD cohort. E. Gene Ontology (GO) and Kyoto Encyclopedia of Genes and Genomes (KEGG) enriched pathway bar chart of Differentially expressed genes (DEGs) in metastasis versus non-metastasis colon cancer. F. Volcano plot showing nearly all the genes enriched in pathway of NK cell-mediated cytotoxicity were downregulated in the metastasis colon cancer. G. Gene set enrichment analysis (GSEA) of KEGG gene set. H. Natural killer cell mediated cytotoxicity was enriched in the non-metastasis colon cancer.

To further understand the potential triggers that induce colon cancer metastasis, the DEGs were calculated between the metastasis and non-metastasis colon cancer groups in the TCGA COAD cohort. A total of 1,378 DEGs were determined using the limma package with cutoffs |log fold change|>1.5 and p <0.05, including 817 upregulated and 561 downregulated genes between metastasis and non-metastasis colon cancer. Subsequently, Gene Ontology (GO) and Kyoto Encyclopedia of Genes and Genomes (KEGG) enrichment analyses were performed to investigate the correlative functions and pathways, As shown in Fig. 3E, cell killing, NK cell-mediated immunity, NK cell-mediated cytotoxicity, NK cell activation, and MHC-related GO terms and the KEGG pathways of NK cell-mediated cytotoxicity were significantly enriched. Interestingly, nearly all the genes enriched in pathway of NK cell-mediated cytotoxicity were markedly downregulated in the metastasis group compared to the non-metastasis group (Fig. 3F). Similarly, Gene Set Enrichment Analysis (GSEA) revealed that NK cell-mediated cytotoxicity was enriched in the non-metastasis group (Fig. 3G and H).

### The Landscape of NK Cells in CCLM Progression

Considering the differences in infiltrated NK cells between tumor-adjacent tissues and tumor tissues as well as colon cancer with and without metastasis, we further focused on NK cells to explore the detailed difference of NK subgroups in the CCLM microenvironments. Unsupervised clustering analysis of NK cells was thus performed and Eight NK cell subtypes were thus identified (Fig. 4A). Combining the highly variable features of each NK cluster (Fig. 4B) and CIBERSORT-reported canonical NK cell markers (Fig. 4C and D), NK cells were classified into three cell types, including activated NK cells identified by the expression of KIR2DL4, GPR183, GRP171, CD69, and IFNG, resting NK cells marked by GZMK, TTC38, CD160, and PLEKNF1, and the other NK cells of which no characteristic gene was identified. Although all three NK cell subtypes were presented in primary colon tumors, adjacent normal colon tissues, liver metastasis tumors, and lymph nodes (Fig. 4E), the infiltration grade for each of these NK cell subsets was significantly different. At the individual sample level, considerable variability was observed in the NK cell subset composition. Where serial samples were available from individual patients, the expression of resting NK cells increased, whereas that of activated NK cells decreased during disease progression (Fig. 4F and G). Additionally, using the paired primary colon tumor, adjacent normal colon tissues, and liver metastasis tumor samples, we observed that the gradient of the resting NK cells increased, whereas the gradient of the activated NK cells decreased (Fig. 4H).

**Figure 4:**
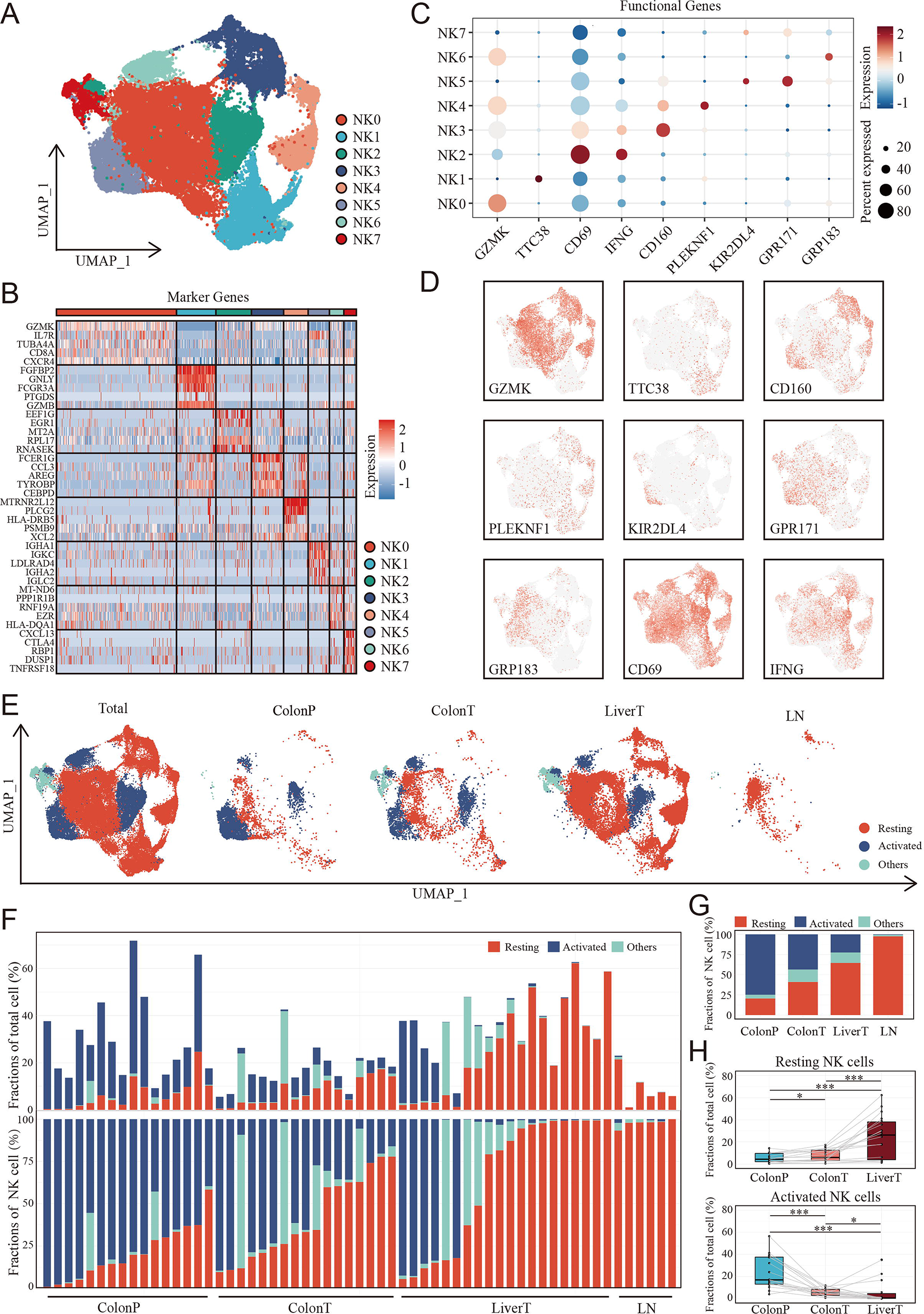
The landscape of NK cells in the disease progression of CCLM. A. The UMAP plot of NK cells from CCLM. B. Unsupervised clustering identifies 8 subsets of NK cells. C. Expression of key markers that distinguish resting and activated subsets of NK cells. D. Expression of key resting and activated NK cell markers across all samples. E. The UMAP plot of distribution of resting and activated NK cells from ColonP, ColonT, LiverT and LN. F. Cellular landscape of each sample from the ColonP, ColonT, LiverT and LN. The proportion of NK cells subsets in total immune cells (upper). The proportion of NK cells subsets in total NK cells (lower). G. Number of cells identified from each group (ColonP, ColonT, LiverT and LN) by cell type proportion. H. Proportions of resting (upper) and activated (lower) subsets of NK cells. P values were determined by the paired nonparameter test.

To evaluate the clinical relationship between NK cell differences and CCLM, CIBERSORT was performed using the TCGA COAD cohort. The percentages of resting NK cells significantly increased (Fig. 5A), and the activated NK cells decreased in the metastasis group (Fig. 5B), which is consistent with the scRNA-seq results. Also, we observed that the resting NK cells were significantly increased, whereas the activated NK cells decreased in higher T, N, and TNM stages (Fig. S5A). In terms of survival, neutrophils and resting NK cells were associated with a worse prognosis, whereas activated NK cells and M1 macrophages were associated with better outcomes (Fig. S5B). Further, survival analysis showed that colon cancer patients with lower and higher degrees of infiltration of activated NK cells and resting NK cells, respectively, had a significantly worse prognosis in the TCGA COAD and GSE29623 cohorts (Fig. 5C and D). Interestingly, resting NK cells were remarkably enriched in the tumor region, whereas activated NK cells were enriched in the non-tumor area of both primary colon cancer and liver metastasis (Fig. 5E).

**Figure.5.**
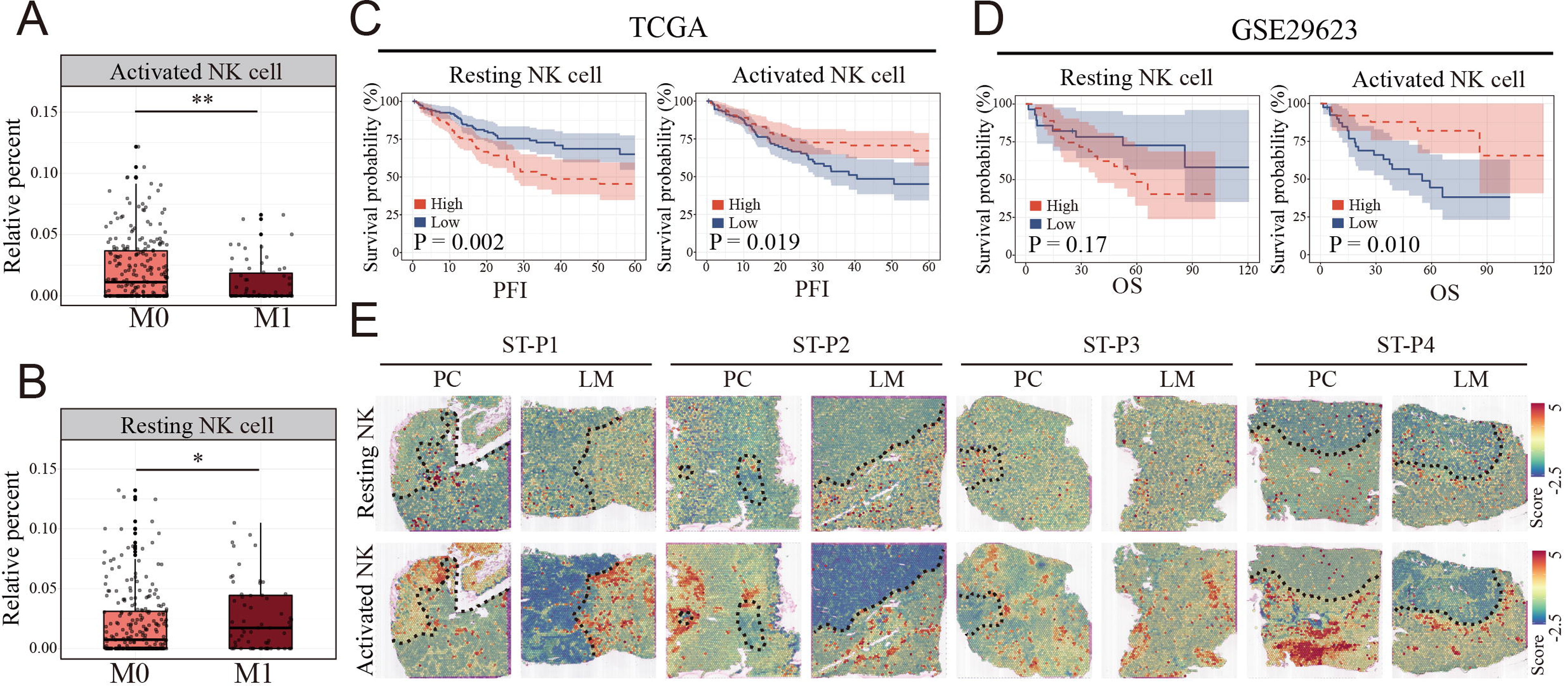
Clinical relationship between NK cells subsets and metastasis of colon cancer revealed by Bulk RNA and spatial transcriptomics. A-B. The relationship of activated and resting NK cell percentage determined by CIBERSORT and tumor metastasis in TCGA COAD cohort. M0: non-metastasis colon cancer; M1: metastasis colon cancer. C-D. K-M survival plots show that high resting NK cell and low activated NK cell predicted poor prognosis in TCGA COAD and GSE29623 cohort. PFI: Progression Free Interval; OS: Overall Survival. E. The signature scores of resting (upper) and activated NK cells (lower) in colon cancer and liver metastasis in the spatial transcriptomic sections.

### Characterization and Developmental Course of Differential Subsets of NK Cells in CCLM

To further study the heterogeneity of NK cells subgroup, the proportions of all eight clusters of NK cells were calculated (Fig. S6). KIR2DL4+ GPR171+ activated NK cells were lessened in proportion with the progression of the disease from normal colon to colon cancer and liver metastasis cancer, whereas GZMK+ resting NK cells increased (Fig. 6A and B), suggesting that these two NK subsets were associated with metastasis. ST analysis also confirmed that there was more infiltration of the KIR2DL4+ GPR171+ activated NK cell subset in the non-tumor area and GZMK+ resting NK cells in the tumor region, both primary colon cancer and liver metastasis (Fig. 6C).

**Figure 6.**
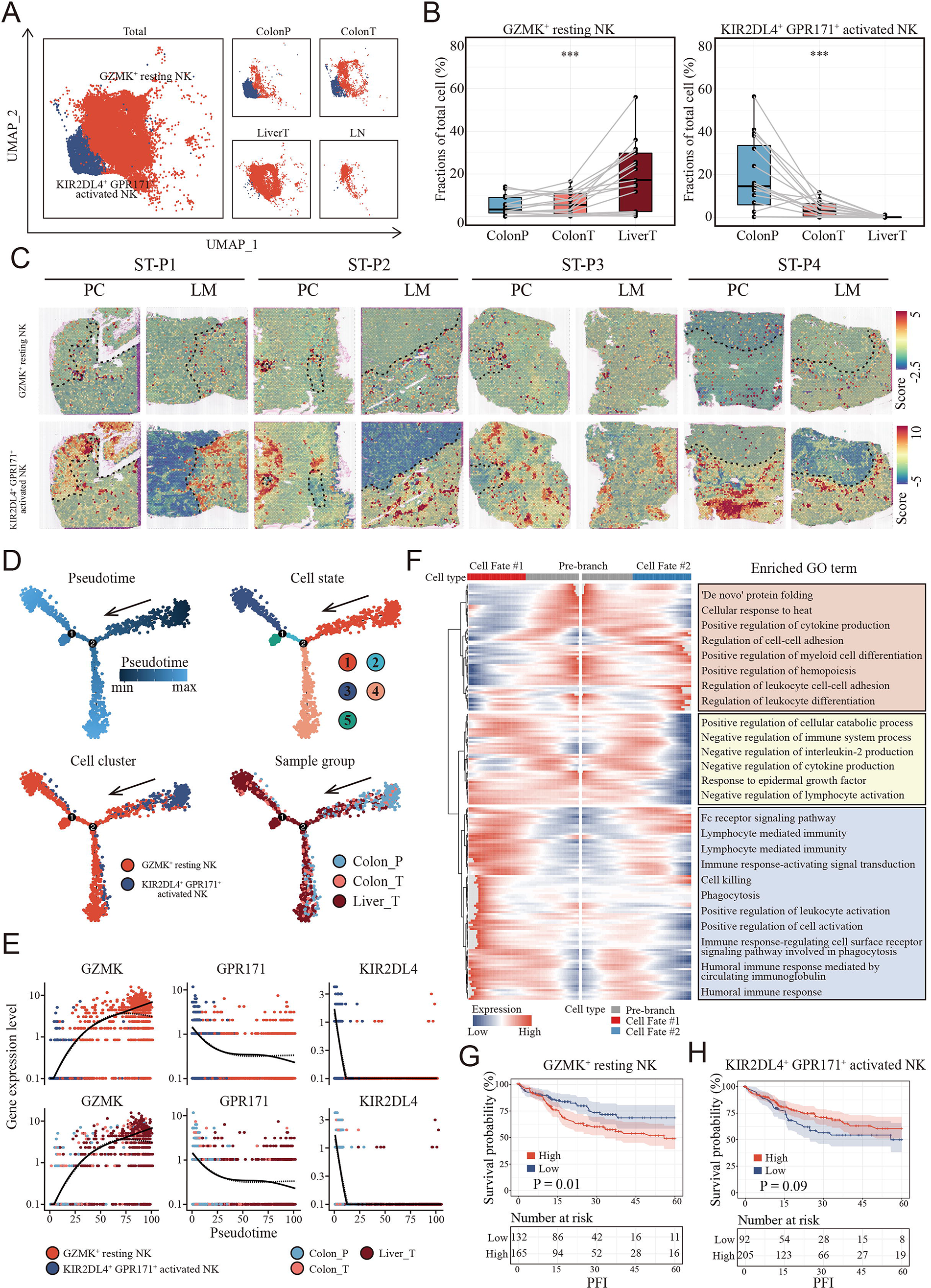
Characterization and developmental course of differential subsets of NK cells in CCLM. A. The UMAP plot of KIR2DL4+ GPR171+ Activated NK cells, GZMK+ resting NK cells from CCLM. B. Proportions of KIR2DL4+ GPR171+ ActiVated NK cells, GZMK+ resting NK cells in total immune cells. P values were determined by the paired nonparameter test. C. The signature scores of GZMK+ resting NK cells (upper) and KIR2DL4+ GPR171+ Activated NK cells (lower) in colon cancer and liver metastasis in the spatial transcriptomic sections. D. Monocle analysis showing the developmental trajectory of NK cells. Color as in Pseudotime, cell state, subsets of NK cells and sample group. E. PseudoLJtemporal change curve of marker genes in each subsets of NK cells. F. The heatmap shows the expression patterns of the top 50 significant genes in branched expression analysis modeling, associated GO terms (using DAVID v6.7) are given on the right of the corresponding gene clusters. G-H. The Kaplan–Meier curve shows COAD patients survival with different GZMK+ resting NK cells and KIR2DL4+ GPR171+ Activated NK cells infiltration.

Pseudotime cell trajectory analysis of the two NK cell clusters was constructed to investigate the evolutionary dynamics of metastasis-associated NK cells. NK cells from the normal colon group were located at the top right corner of the trajectory curve, suggesting the clear starting point of this evolving trajectory curve. After confirmation of this starting point, developmental routes were determined to begin with the KIR2DL4+ GPR171+ activated NK cells and then develop into GZMK+ resting NK cells (Fig. 6D).

To explore the transitional relationships of marker genes in NK cells in distinct clusters, we examined the expression level of these markers in different clusters through pseudotime analysis. Marker genes of activated NK cells (KIR2DL4 and GPR171) exhibited a gradient descent expression pattern, whereas the expression of resting NK cell marker genes (GZMK) exhibited a gradient rise pattern (Fig. 6E). BEAM analysis was performed to identify the cell cluster marker genes that change as NK cell subtypes move from the activated stage to the resting stage. The branched heatmap showed the expression dynamics of the top 200 significant genes in different cell fate branches. Corresponding GO enrichment analyses of these significant genes further demonstrated that a mass of leukocyte differentiation and immune system process-related GO terms were significantly enriched (Fig. 6F).

Interestingly, patients with colon cancer with higher infiltration of GZMK+ resting NK cells (Fig. 6G) and lower infiltration of KIR2DL4+ GPR171+ activated NK cells (Fig. 6H) exhibited shorter survival in the TCGA COAD cohort.

### Tumor Cells-Educated NK Cells Shift Toward Tumor-Promoting Status Depends on Cell-to-Cell Interaction

To verify the NK cell-mediated inductive effect on colon cells, we set up mixed cell co-culture experiments using colon cell line HCT-116 and NK cell line NK-92 in the ratio 1:1 (CN group). As controls, NK cells were cultured alone, either in the tumor supernatant (SN group) or fresh medium (MN group). The co-cultured supernatant of each group were collected to incubate fresh HCT-116 after 24 h of co-culturing. Interestingly, CCK-8 and colony formation assays showed that colon cells in CNS group, which colon cells were cultured in the supernatant of co-cultured system that NK and cancer cell were in contact, underwent a significant increase in proliferation (Fig. 7A-C) compared to the CS group, where cancer cells were cultured directly in the supernatant of cancer cell after 24h culturing. Similarly, the migration and invasion of HCT-116 were also significantly increased in CNS group (Fig. 7D-F).

Furthermore, colony formation assays (Fig. 7G-H), CCK-8 (Fig. 7I), and transwell assays (Fig. 7J-L) showed colon cells in CNS group underwent a significant increase in proliferation, invasion and migration, compared to those cultured in the co-cultured supernatant (SNS group) and fresh medium (MNS group).

**Figure.7.**
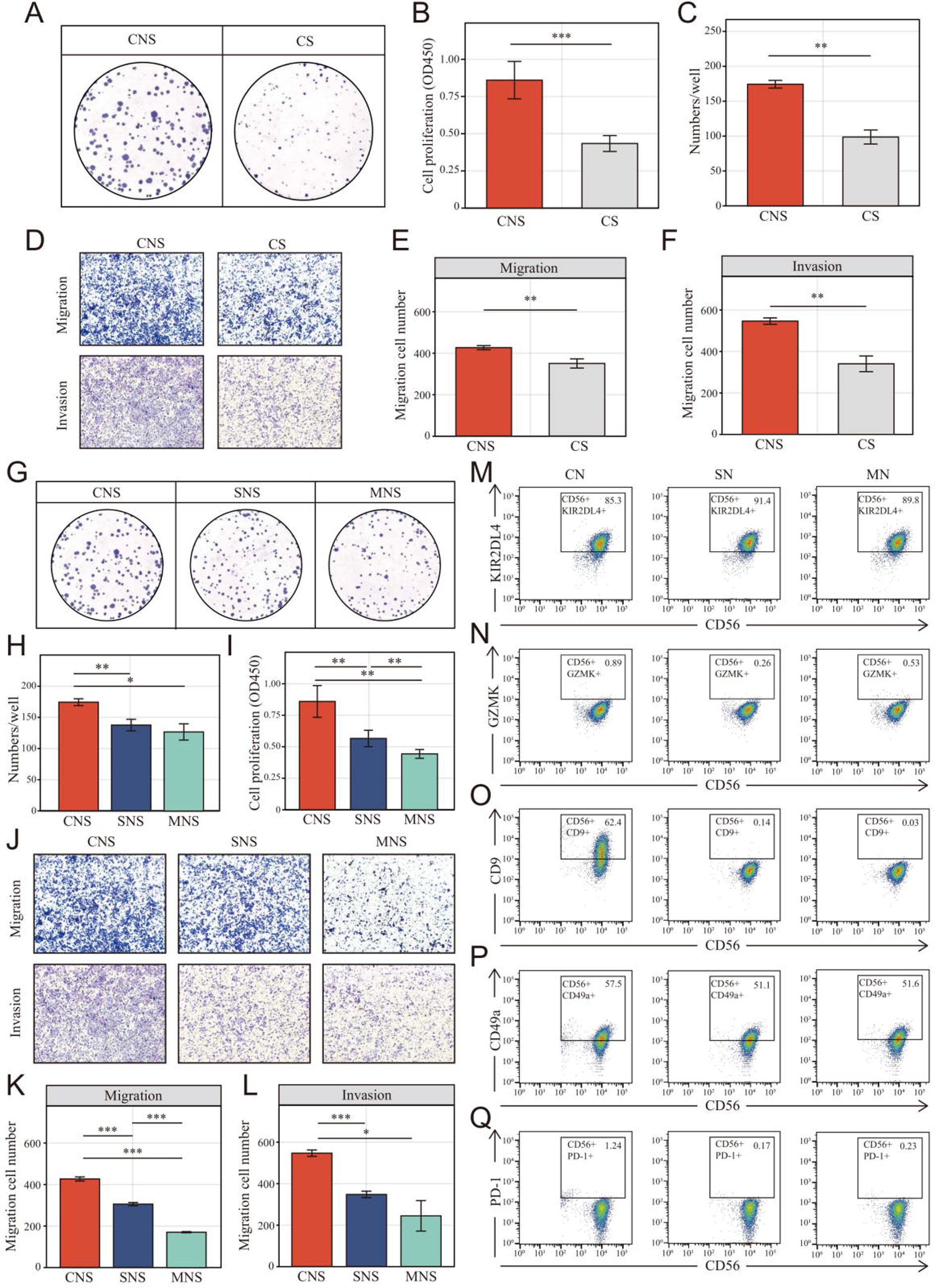
Colon cancer cells (HCT-116) educated NK cells shift toward tumor promoting status depends on cell-to-cell interaction. A-B. Clone formation assay showed the NK cell-mediated inductive effect on cell proliferation of colon cancer cell (HCT-116). Colon cancer cells were cultured in the supernatant from different co-culture system in transwell devices. CNS: colon cancer cells were cultured in the supernatant from co-culture system that NK cells and cancer cells were cultured in direct contact (CNS); CS: cancer cells cultured directly in supernatant of cancer cells. C. Cell Counting Kit-8 (CCK-8) assay showed the NK cell–mediated inductive effect on cell proliferation of colon cancer cell. D-F: The NK cell–mediated inductive effect on migration and invasion of colon cancer cell. G-H: Clone formation assay CCK-8 assay showed the NK cell–mediated inductive effect on cell proliferation of colon cancer cell. SNS: colon cancer cells were cultured in the supernatant from co-culture system that NK cells were cultured in supernatant of cancer cells; MNS: colon cancer cells were cultured in the supernatant from co-culture system that NK cells were cultured in fresh medium. M-Q. Phenotype switch (KIR2DL4, GZMK, CD9, CD49a, PD-1) of NK cells was induced by cell-to-cell interactions with cancer cells. CN: NK cells were co-cultured with colon cancer cells; SN: NK cells were cultured in supernatant of cancer cells; MN: NK cells were cultured in fresh medium.

Additionally, another colon cancer cell line, DLD-1 was chosen to evaluate the effect of the supernatant in the different co-culture groups. DLD-1 cells in the CNS group did not undergo a prominent increase in proliferation compared to CS group (Fig. S7A-C). The same results were obtained for their migration and invasion (Fig. S7D-F). Similarly, no such cell functional changes were induced in the CNS group compared to SNS and MNS groups (Fig. S7G-L).

Given the characteristic spatial distribution of NK cell subpopulations, we hypothesized that the tumor region-enriched NK cells might be educated by tumor cells and thus functionally distinct. To verify the possible effect of tumor cells on the phenotypic switch of NK cells, NK cells were analyzed using FACS to determine the expression of the NK cells markers above identified (GZMK and KIR2DL4) as well as tumor infiltrated NK cells markers: CD9, CD49a and PD-1. FMO control was set up to define the position of the positive gate as shown in Fig.S8. Upon exposure to tumor cells, NK cells significantly decreased and slightly increased the expression of activated marker KIR2DL4 (Fig. 7M) and the resting marker GZMK (Fig. 7N), respectively. On the other way, for GZMK, which known as the secretory protein, flow cytometric analyses were also performed with cell fixation and permeabilization. However, nearly all the NK cells in different groups were GZMK positive and no significant differences were found among each group (Fig.S9). Additionally, NK cells in CN group underwent an increasing of tumor infiltrated NK cells markers, CD9 (Fig. 7O), CD49a (Fig. 7P) and PD-1(Fig. 7Q), especially CD9. However, no such effect was observed after co-culturing with tumor supernatant and fresh medium.

On the other way, NK cells were also co-cultured with HFF, a normal human cell line. The tissue infiltrated NK phenotype identified above (CD9, CD49a, PD-1) were determined. When co-cultured with HFF in direct contact (CN group), NK cells were also tending towards tissue infiltration state. However, the domestication effect was significantly reduced compared to co-culturing with tumor cells (Fig.S10A and B). Additionally, fresh HCT underwent a limited increase (no statistical significance was found) in migration when cultured in CNS group, but not in the SNS group MNS group (Fig.S10C-E).

### Resting NK-induced Colon Cancer Malignant Phenotype Promotion Depends on SCF Release

To study the potential mechanism by which the supernatant from the NK and colon cells in direct contact with the co-culture system promoted colon cell migration, Luminex liquid suspension chip detection was used to compare the differential expression of 48 common chemotactic and inflammatory cytokines in the CNS, SNS, and MNS groups. We observed that SCF was upregulated in all three repetitions in the CNS group (Fig. 8A-B). Further, ELISA confirmed the upregulated level of SCF in the CNS group (Fig. 8C).

**Figure.8.**
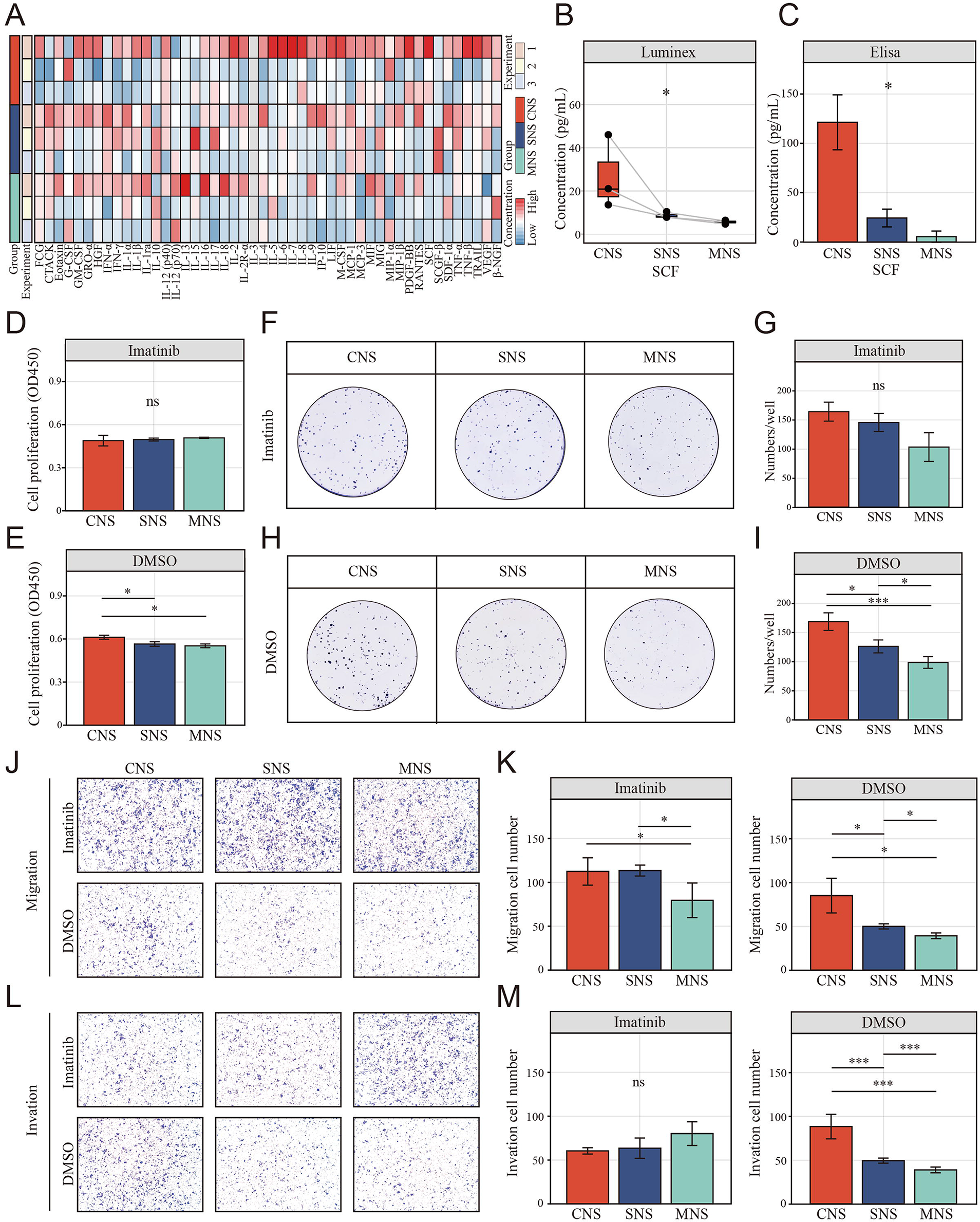
The resting NK cell promote tumor malignant phenotype via elevating tumor-derived sSCF. A. luminex liquid suspension chip detection of 48 common chemotactic and inflammatory cytokines in CNS, SNS and MNS group. B-C. Concentration of SCF determined by luminex liquid suspension chip and Elisa in CNS, SNS and MNS group. D-E. CCK-8 assay showed the proliferation of HCT-116 cells was inhibited by imatinib mesylate, evaluated by a CCK-8 assay. Cells were incubated in the supernatant from different co-culture system with DMSO or 2 µM imatinib mesylate. F-I. Clone formation assay showed the proliferation of HCT-116 cells was inhibited by imatinib mesylate. J-M. The NK cell–mediated inductive effect on migration and invasion of HCT-116 was inhibited by imatinib mesylate.

We further assessed the inhibitory effects of imatinib mesylate on the enhanced proliferation and invasion. CCK-8 (Fig. 8D-E) and colony formation assays (Fig. 8F-I) showed that imatinib mesylate (2 µM) significantly inhibited supernatant-enhanced proliferation of HCT-116. Similarly, the supernatant-enhance migration (Fig. 8J-K) and invasion (Fig. 8L-M) were also inhibited by imatinib.

## Discussion

CCLM is a multistep process and is mostly fatal(44). During disease progression, functional interactions between tumor cells and the surrounding TME are critical for influencing tumor growth, promoting angiogenesis, and finally resulting in metastasis(45). Since the genomic divergence between primary and metastatic CRC cells is relatively low(46, 47) and accumulating evidence has highlighted the key role of tumor-infiltrating immune cells in dictating the fate of cancer cells(48, 49), we presented a comprehensive cellular and spatial immune landscape of the primary and liver metastatic tumors of CRC by using previously published scRNA-seq and ST data. Consistent with previous studies(30, 50), we observed that the immune microenvironment of primary and liver metastasis lesions of colon cancer had undergone extensive remodeling with a significantly decreased proportion of NK cells and a strong enrichment of T cells. However, the contrast between the decreased proportion of NK cells in tumor tissue and the enrichment of NK cells in tumor regions may indicate the specific effects of NK cells in CCLM.

Traditionally, NK cells within the TME were known as potential suppressors for tumor growth by directly killing cells and secreting proinflammatory cytokines(51, 52). However, the function of NK cells highly depends on their maturation status and localization(53). The peripheral blood CD56^dim^CD16^high^ NK cell population predominantly mediates the killing of target cells by secreting perforin and granzymes(54), whereas the CD56^bright^CD16^low^ NK cells that reside in secondary lymphoid tissues are immature and have reduced cytotoxic capability(55). Recently, tumor-associated NK cells, which are enriched in tumors with impaired anti-tumor functions, were discovered to be associated with an unfavorable prognosis and resistance to immunotherapy in multiple solid tumors (56), such as lung cancer, pancreatic cancer, and esophageal cancer. The potential impact of NK cells on CCML progression is still poorly investigated.

In this study, the proportion of NK cells was higher than in past cognition, similar to a previous study(57) in that NK cells occupied the largest immune cell compartment of nearly 50% in cholangiocarcinoma. In contrast, NK cells may not be well distinguished during automatic annotation using SingleR. Consistent with the study that revealed that treatment responders are associated with TME remodeling, including NK cell recruitment(58), the proportions of NK cells in PR patients were significantly higher among the LiverP and LiverT groups (Fig. S2D and E), which also increased the overall proportion of NK cells to some extent. Additionally, we observed that NK cells are decreased in primary colon cancer and liver metastatic using scRNA-seq, consistent with previous findings supporting that NK cells play a crucial role in anticancer immunity(59, 60). However, we further subdivided NK cells considering the conflicting result, in which NK cells were significantly enriched in the tumor area when using ST. Since this is the first study to comprehensively explore the heterogeneity of NK cells, no previous NK cell subpopulation markers were available; resting and activated NK cell markers in CIBERSORT were thus used to annotate NK cells. Consistent with a recent study that reported that mature and immature NK cells serve different functions(61), our results also provide evidence that resting NK cells in the colon cancer TME possess a potential tumor-promoting activity, whereas activated NK cells play an anti-tumor role as conventional cognition. Bulk RNA-sequencing and the corresponding clinical data further indicated that higher infiltration of resting NK cells and lower infiltration of activated NK cells were correlated with a worse prognosis in patients with CRC. However, since the resting and activated NK cells we identified were only the aggregation of multiple NK cell subpopulations, further studies are required to investigate the key NK cell subpopulations that play a crucial role in CCLM.

Further analysis of NK cell subtypes revealed a dramatic change in KIR2DL4+ GPR171+ activated NK cells and GZMK+ resting NK cells, whereas no other NK cell subpopulations differed in colon cancer progression. KIR2DL4, also known as CD158d, is an unusual member of the killer cell Ig-like receptor family expressed in all NK cells(62). Studies revealed that KIR2DL4 activates the cytotoxicity of NK cells(63), and NK-92 cells stimulated with KIR2DL4 had higher cytotoxicity against breast cancer cells(64). Consistent with previous studies, we also observed that the expression of the KIR2DL4+ GPR171+ activated NK cell subtype was decreased during CCLM progression, possibly reflecting the anti-tumor activity of this population. However, the GZMK+ resting NK cell subsets were first identified as potential tumor promoters. Moreover, it has been proven that GZMK^high^ CD8^+^ T effector memory cells, a cell subset that is particularly similar to GZMK+ NK cells, were associated with poor clinical outcomes in patients with colorectal tumors. Our results also showed that GZMK+ resting NK cells highly infiltrate the cancer region, and this predicts a worse prognosis in patients with colon cancer. However, despite the interesting and gratifying clinical relevance, the mechanism remains unknown.

Considering the evolutionary trajectory and the characteristic distribution of these metastasis-associated NK cell subtypes, we predicted that activated NK cells might differentiate into resting NK cells under the action of tumor cells. Consistent with a recent study that indicated that exposure to cancer cells causes NK cells to lose their cytotoxic ability(28), we identified the tendency of NK cells to differentiate to an resting state with decreasing expression of activated marker KIR2DL4 as well as increasing expression of decidual-like NK cells markers: CD9, CD49a and PD-1(27), which are only triggered by cell-to-cell interactions as validated by FACS. It is worth mentioning that decidual-like NK cells were the newly identified subset of NK cells that promote tumors(27).

Additionally, NK-mediated tumor cell editing appears to partly depend on the release of cytokines in in vitro co-culture experiments. SCF, the natural ligand of c-Kit with membrane-bound (mSCF) and soluble forms (sSCF)(65), enhances the growth and migration of cancer cells such as cervical cancer(66), Ewing’s sarcoma(67) and CRC(43). Although the dimeric mSCF is remarkably more active and induces a more persistent activation of the receptor(65, 68), the tumor-promoting effect of the supernatant from the co-culture system in the current study and the reversion effect of imatinib for the co-culture supernatant induced pro-oncogenic effects suggests an important role of sSCF on colon cancer cells. However, no up-regulation of KITLG (the SCF encoding gene) was found in HCT-116 after co-culturing with NK cells (Fig. S11A). In addition, when using DLD-1, a colon cancer cell with a nearly 15 times lower expression of KITLG than HCT-116 (Fig. S11B), we also did not observe co-cultured supernatant induced tumor-promoting effects. We thus hypothesized that in our experimental system, NK cells may promote the cleavage and release of the extracellular part of SCF on the tumor cell surface after direct contact with tumor cells, and thus increase the concentration of sSCF, which induces the malignant phenotype of colon cancer cells in a paracrine manner (Fig. 9). However, the detailed mechanism still needs to be further studied.

**Figure.9.**
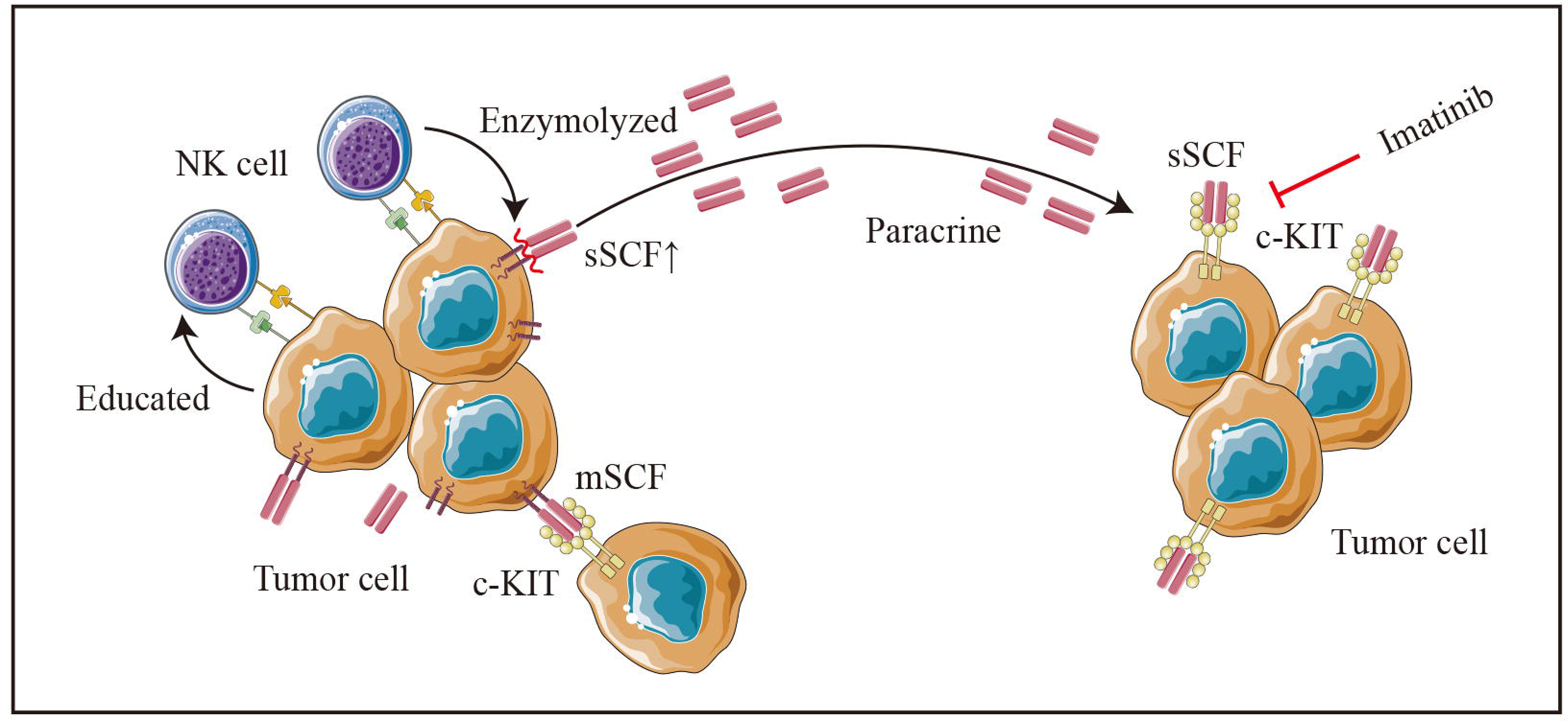
Schematic diagram. Colon cancer cells (HCT-116 cells) educate NK cells as resting status depends on cell-to-cell interaction. Tumor-educated NK cells subsequent enhances tumor malignancy in a paracrine manner by elevating tumor-derived sSCF.

In summary, we have unveiled the spatial and cellular profiles of TME from tumors and paratumor tissues of CCLM. Our analysis uncovered the different states, functions, and dynamic nature of NK cells in different CCLM settings, which can be used for further identification of regulatory mechanisms and for developing potential therapeutic targets.

## Data Availability

All data generated or analyzed during this study are available from the corresponding author on reasonable request.

## Funding

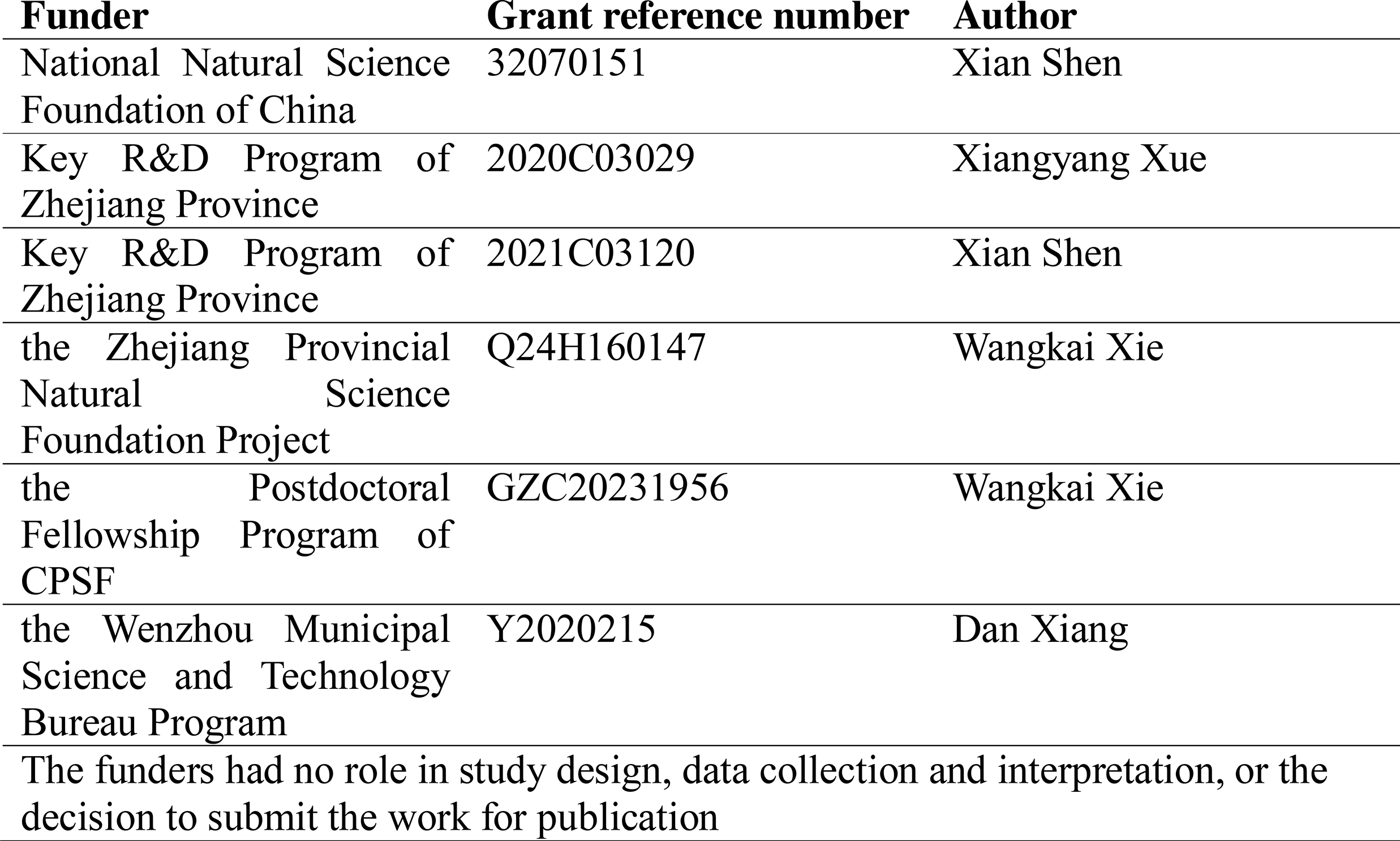

## Acknowledgement

We thank Prof. Qiang Gao from Fudan University and Prof. Xiaoming Zhang from Shanghai Institute of Immunity and Infection, CAS for sharing the SC RNA sequencing and ST data. We thank Zhejiang Key Laboratory of New Techniques for Diagnosis and Treatment of Critical Diseases of Pancreas and Liver and Zhejiang International science and Technology Cooperation base for tumor transformation research for supporting this study. We thank Editage Group (https://www.editage.cn/) for polishing the draft of this manuscript.

]

## Authors’ Contributions

**CCM** and **YYC**: Conceptualization, experimental design and execution, data curation, method development, formal analysis, statistics, writing original draft, and editing the manuscript. **DX** and **TMZ**: conceptualization, data curation and method development. **YXL**, **CL** and **JYY**: experimental design. **MHL** and **TM**: experimental design. **WKX, DFM** and **ZH**: Validation. **DX**, **MDL** and **XS**: Experimental design, supervision, funding acquisition, and manuscript editing. **XYX XS**: Conceptualization, supervision, writing manuscript, project administration.

## Supplemental Figures legends

**Figure S1. Schematic overview of the in vitro experimental design.** A. Schematic overview of coculture experiments design. B. Schematic overview of coculture experiments design in transwells.

**Figure S2. Cluster characterization of the global landscape of CCLM revealed by single-cell transcriptomics.** A. The UMAP plot of all 12 cell clusters. B. Dot plots showing average expression of marker genes in indicated cell clusters. C. The pie chart of the proportion of main immune cells across tissues. D. The proportion of NK cells in total immune cells of each sample (including PR, PD/SD and untreated patients) from the Liver P, Liver T, Colon P, Colon T. E. Proportions of NK cells in PR, PD/SD and untreated patients among Liver P, Liver T, Colon P, Colon T groups.

**Figure S3. Cellular identification in spatial transcriptomic samples.** A. The UMAP plot of all 11 cell clusters. B. Dot plots showing average expression of marker genes in indicated cell clusters. C. Expression of key markers across all samples.

**Figure S4. Gene expression features of each sample. The feature plots showed the expression distributions of ALB, CD24, COL1A1, MYH11, CXCL8, CD14, CD79A AND CD3D in each spatial transcriptomic section.**

**Figure.S5 Clinical relationship between NK cells subsets and colon cancer revealed by TCGA COAD cohort.** A. The relationship of activated and resting NK cell and clinical characteristics of colon cancer in TCGA COAD cohort. B. The relationship of 22 immune cells percentage determined by CIBERSORT and prognosis of colon cancer in TCGA COAD cohort.

**Figure.S6 The proportions of eight clusters of NK cells in Colon P, Colon T, Liver T and LN.**

**Figure.S7 NK cell-mediated tumor promoting effect in colon cancer cells (DLD-1).** A. Clone formation assay showed the NK cell–mediated inductive effect on cell proliferation of DLD-1 cell between CNS and CS groups. B-C: CCK-8 assay showed the NK cell–mediated inductive effect on cell proliferation of DLD-1 cell between CNS and CS groups. D-F. The NK cell–mediated inductive effect on migration and invasion of DLD-1 cell between CNS and CS groups. G. CCK-8 assay showed the NK cell-mediated inductive effect on cell proliferation of DLD-1 cell among CNS, SNS, MNS groups. H-I. Clone formation assay showed the NK cell-mediated inductive effect on cell proliferation of DLD-1 cell among CNS, SNS, MNS groups. J-L The NK cell–mediated inductive effect on migration and invasion of DLD-1cell among CNS, SNS, MNS groups.

**Figure.S8 The positive gate locations of CD56, GZMK, KIR2DL4, CD9, CD49a, PD-1 defined according to the FMO control.**

**Figure.S9 Phenotype switch (CD56+, GZMK+) of NK cells was analyzed by FACS after fixation and permeabilization in different co-cultured groups.**

**Figure.S10 Phenotype switch of NK cells in different co-cultured system and the corresponding NK cell-mediated effect on cell migration of fresh colon cancer cell (HCT-116).** A-B: NK cells underwent phenotype switch (high expression of CD9) when cocultured with HCT and HFF, the phenotype switch was more obvious when co-cultured with HCT. CN: NK cells cocultured with HCT/HFF; SN: NK cells cocultured with supernatant of HCT/HFF; MN: NK cells cocultured in fresh medium. C-E: Transwell assay showed the only tumor co-cultured NK mediated the inductive effect on cell migration of colon cancer cell (HCT-116). CNS: Colon cancer cells were cultured in the supernatant from co-culture system that NK and HCT/HFF were cultured in direct contact; SNS: Colon cancer cells were cultured in the supernatant from co-culture system that NK cocultured with supernatant of HCT/HFF; MNS: Colon cancer cells were cultured in the fresh medium.

**Figure.S11 Relative KITLG expression in different groups.** A. RT-qPCR analysis of KITLG expression in HCT-116 with and without co-cultured with NK cells. B. RT-qPCR analysis of KITLG expression in HCT-116 and DLD-1 cells.

