## Supplementary figures and images for "Resting natural killer cells promote the progress of colon cancer liver metastasis by elevating tumor-derived sSCF"

### Figure-S1

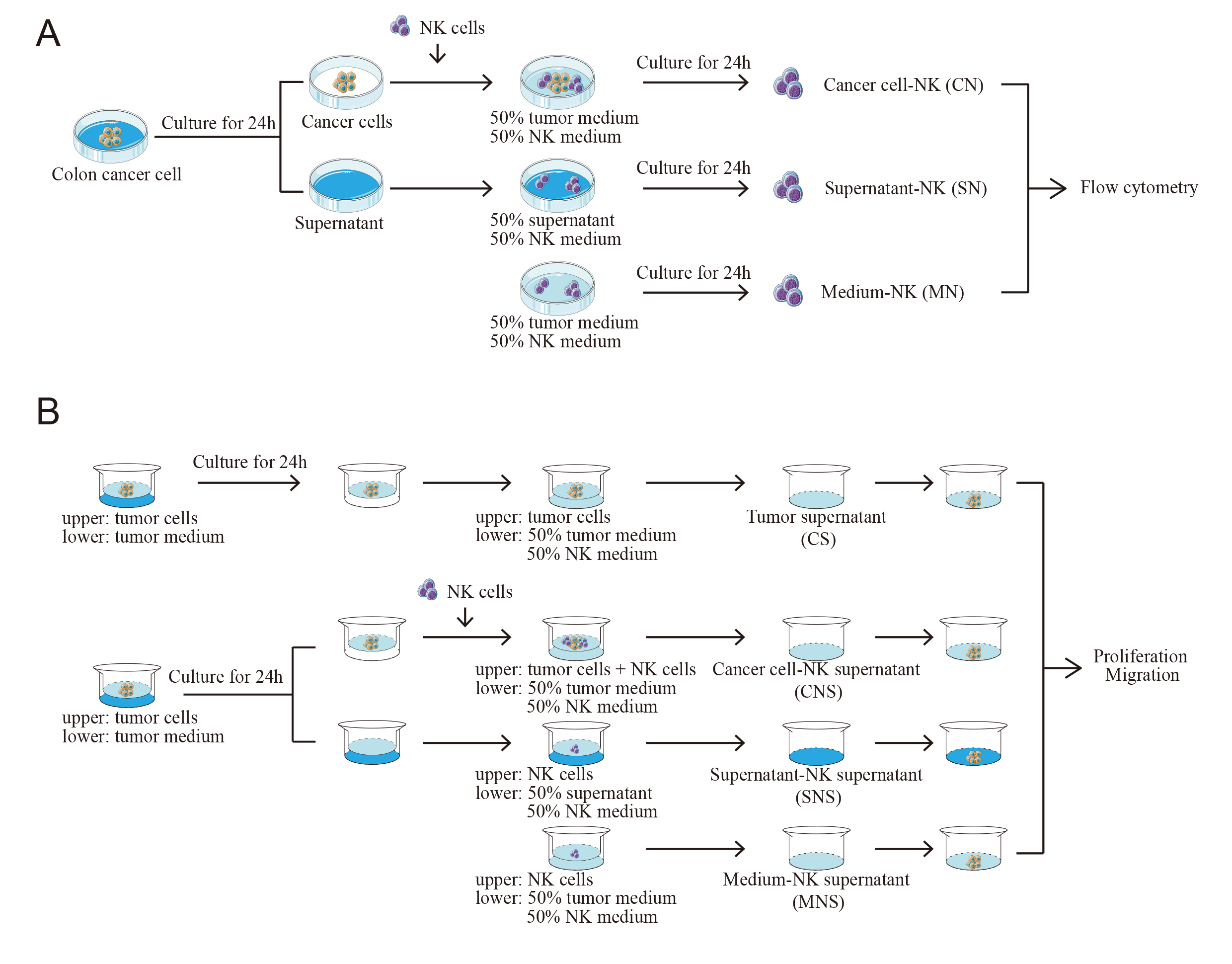

### Figure-S2

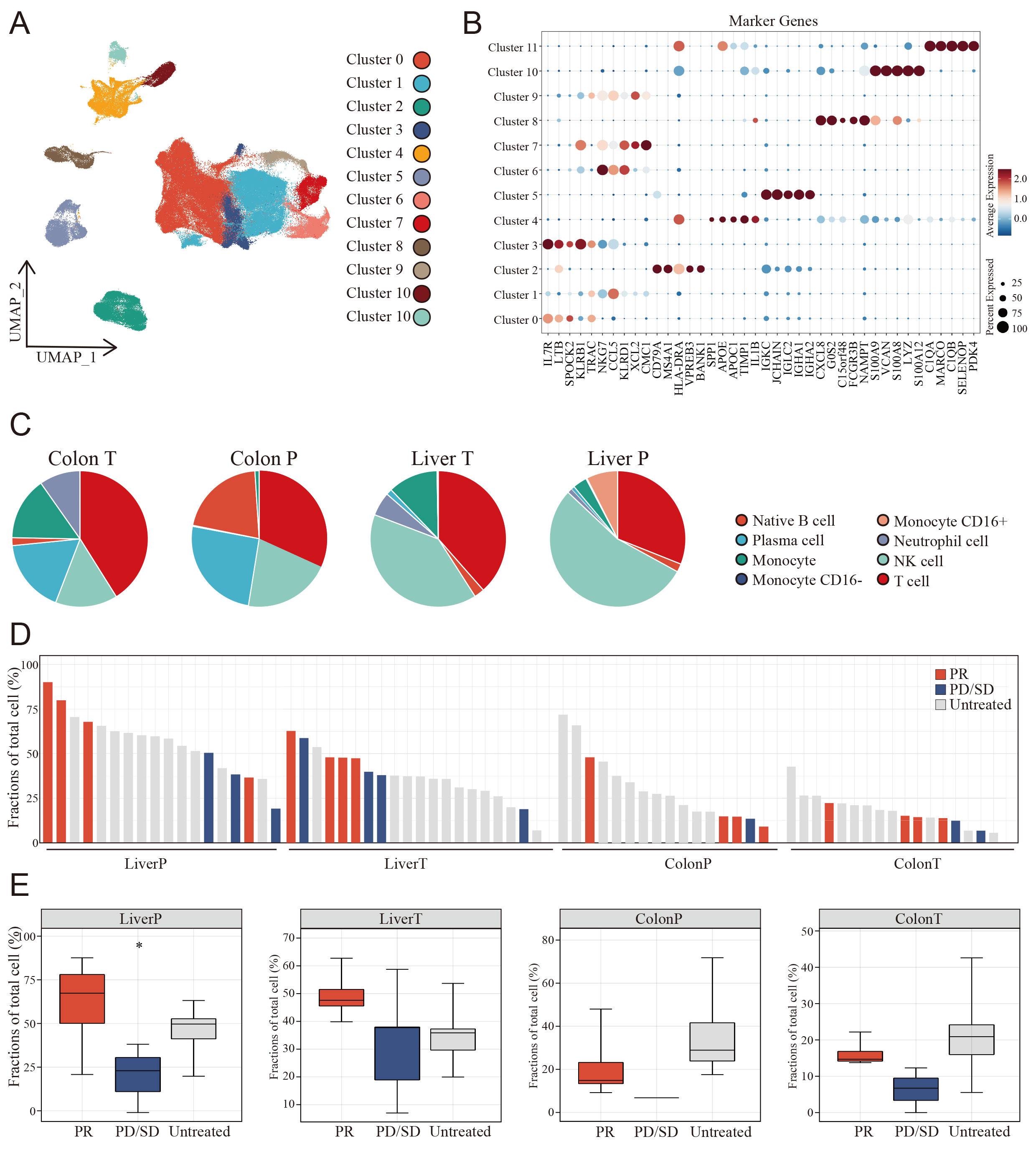

### Figure-S3

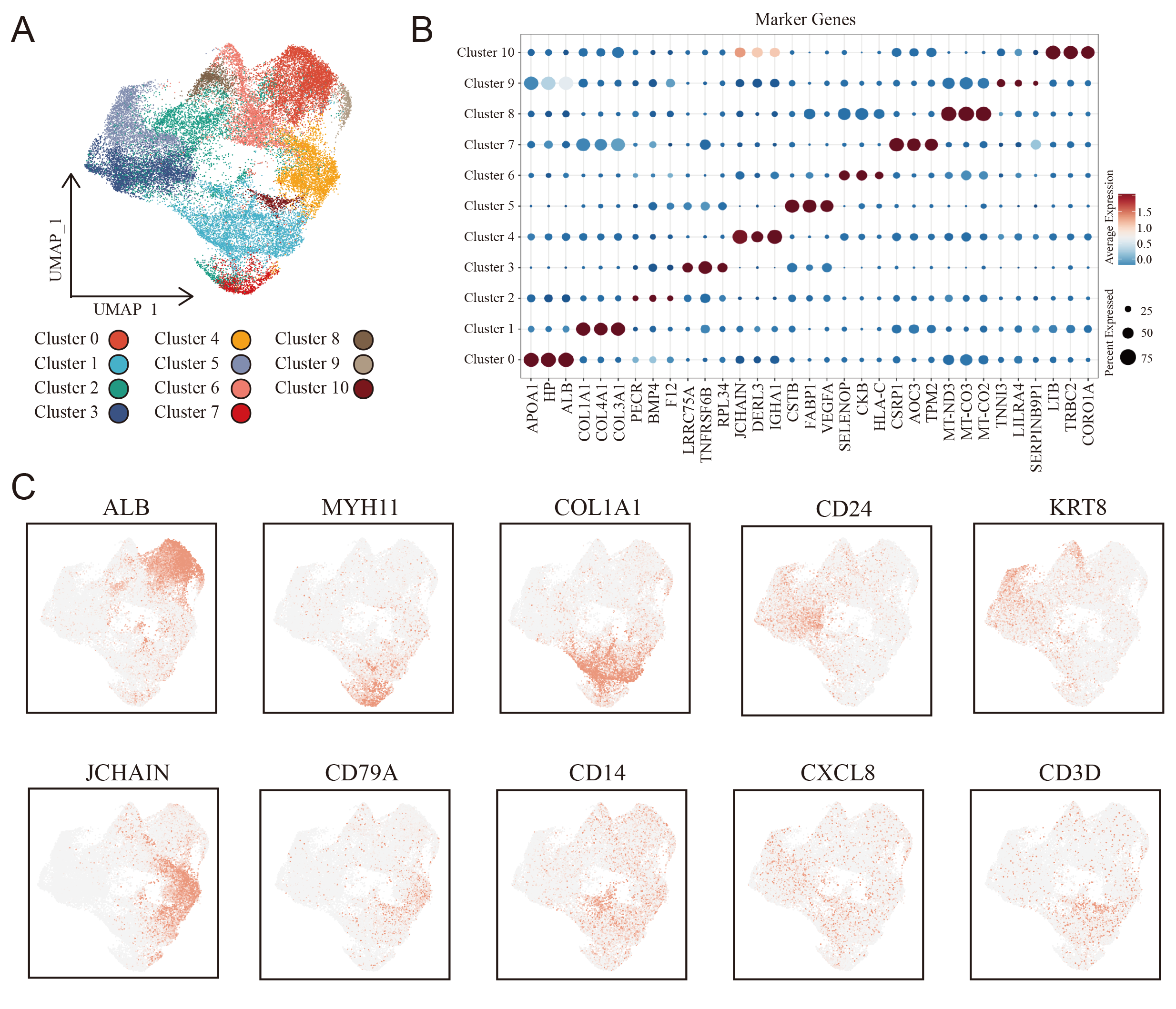

### Figure-S4

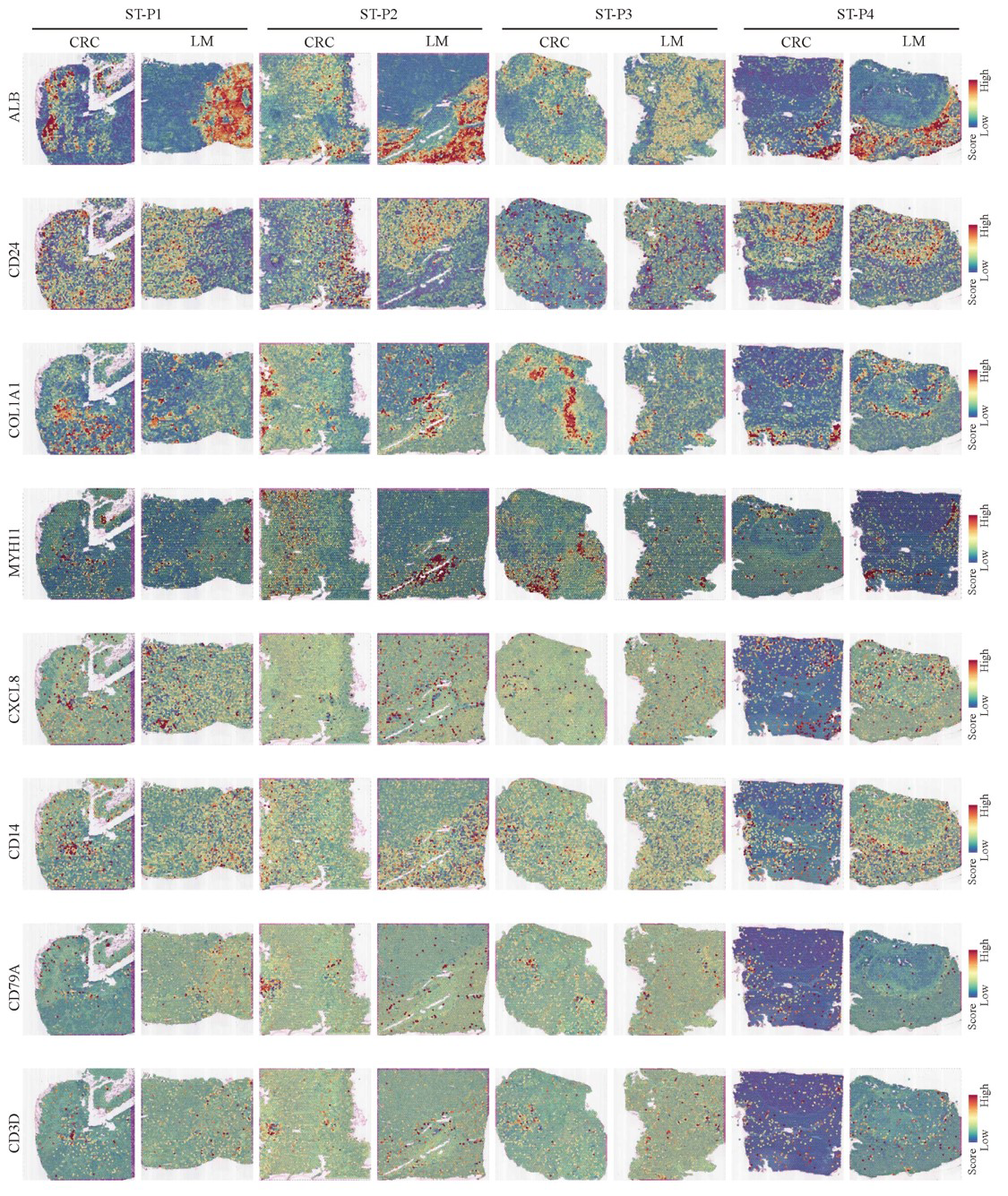

### Figure-S5

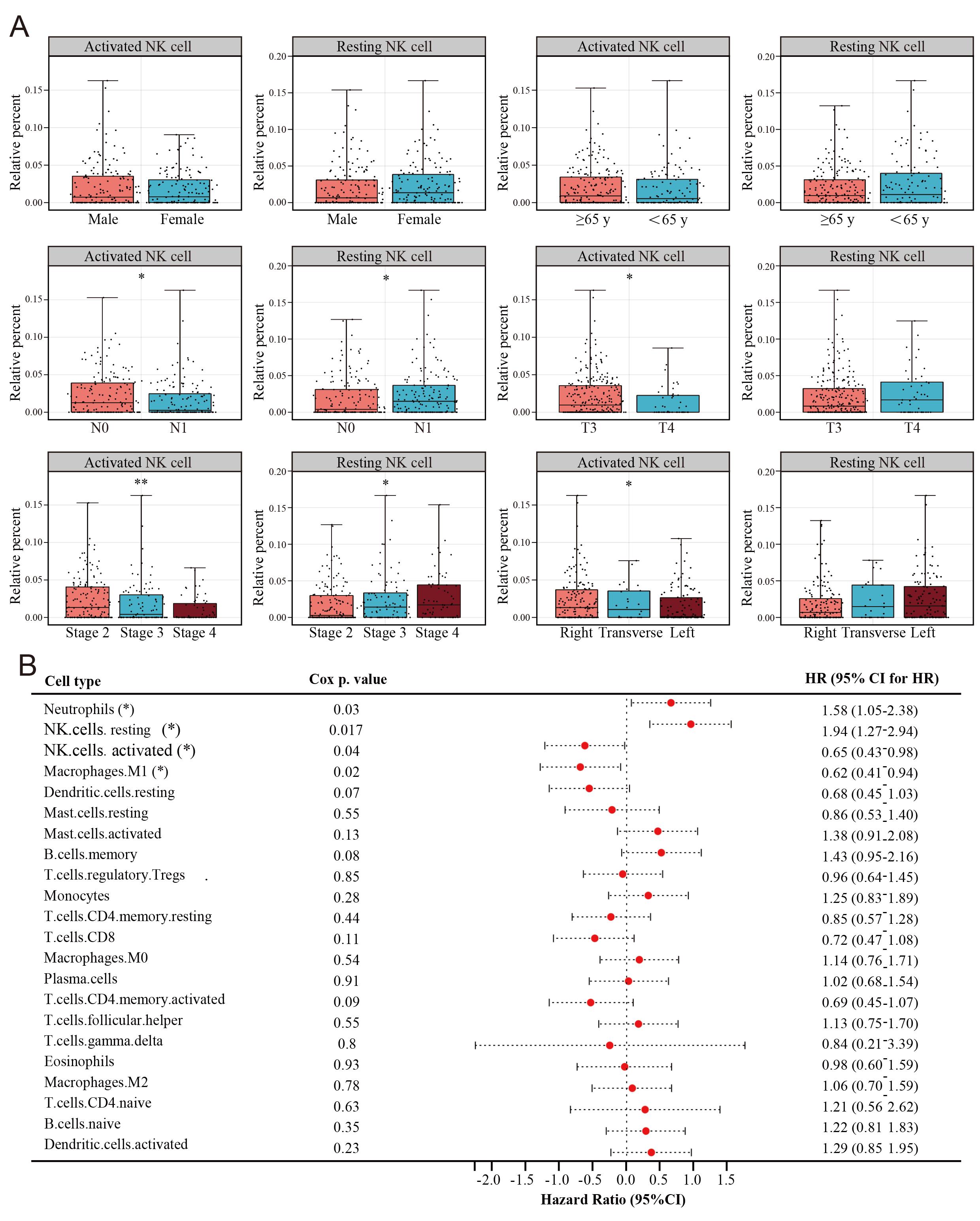

### Figure-S6

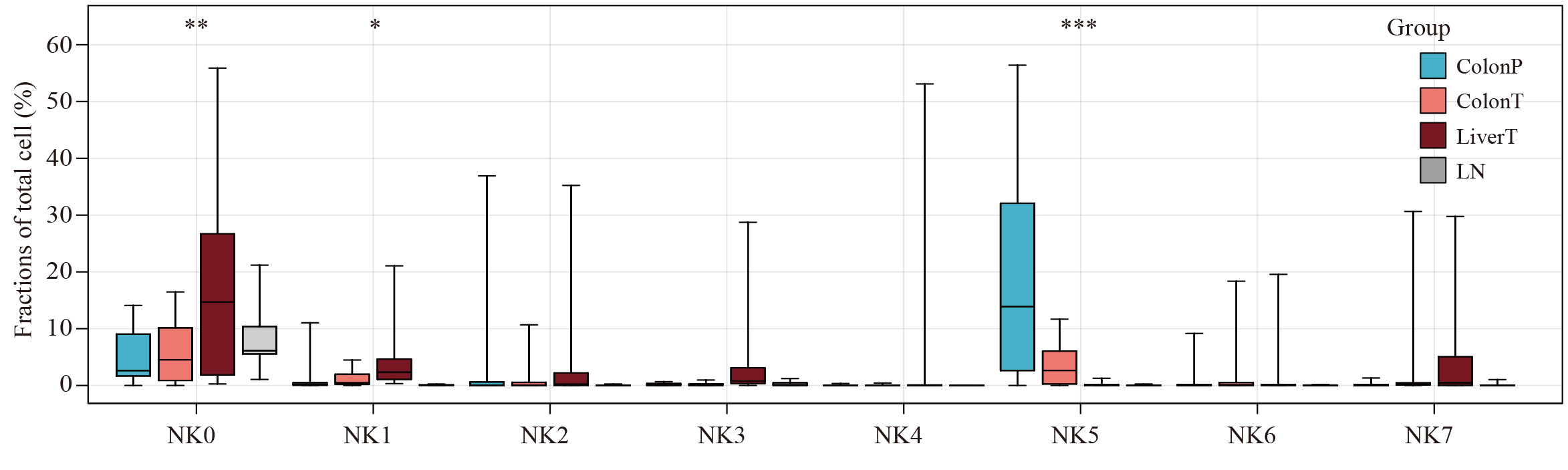

### Figure-S7

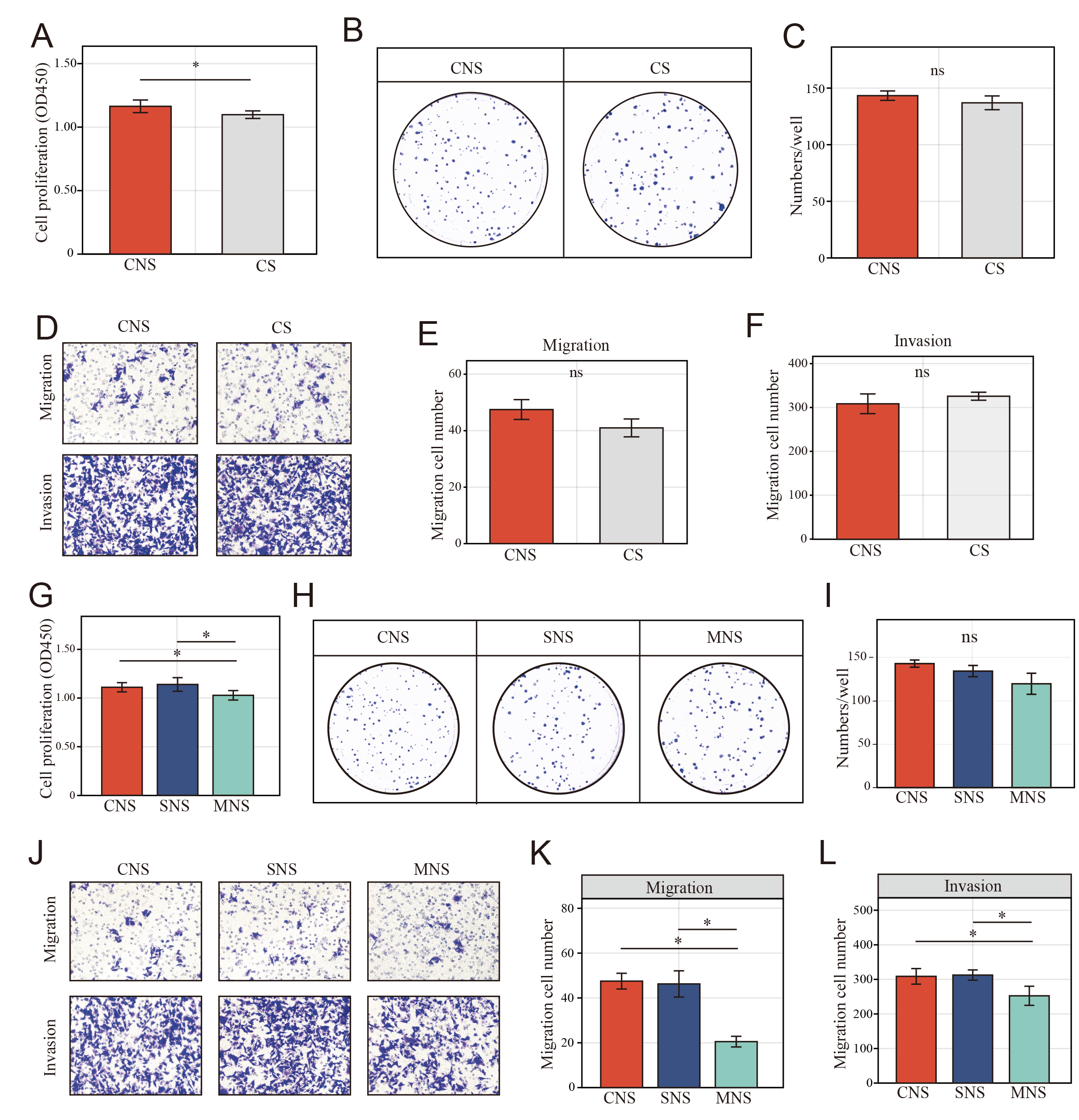

### Figure-S8

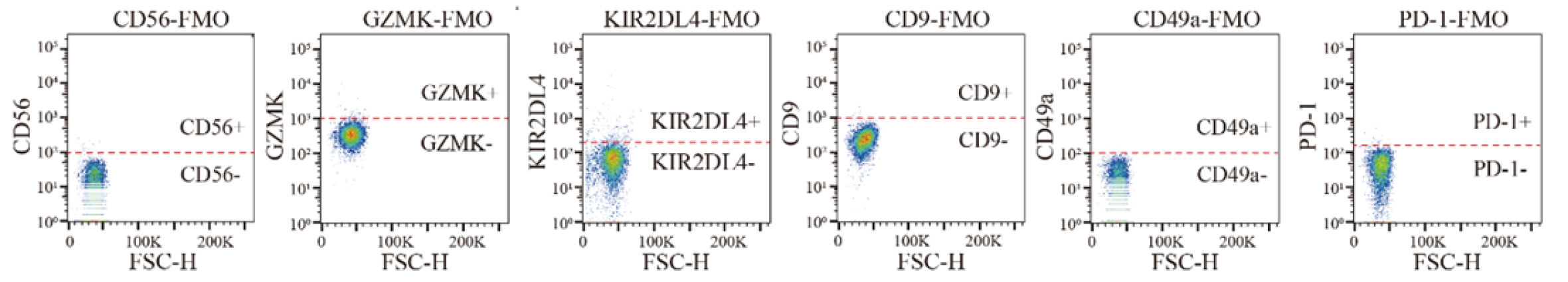

### Figure-S9

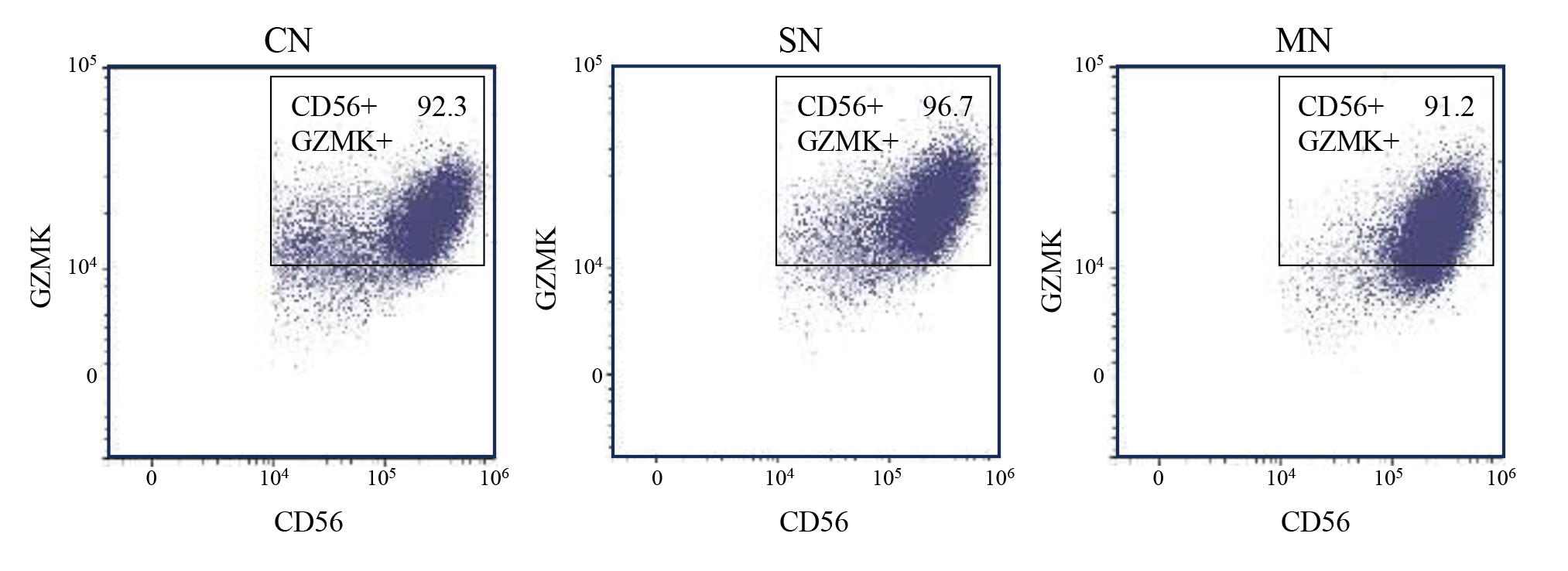

### Figure-S10

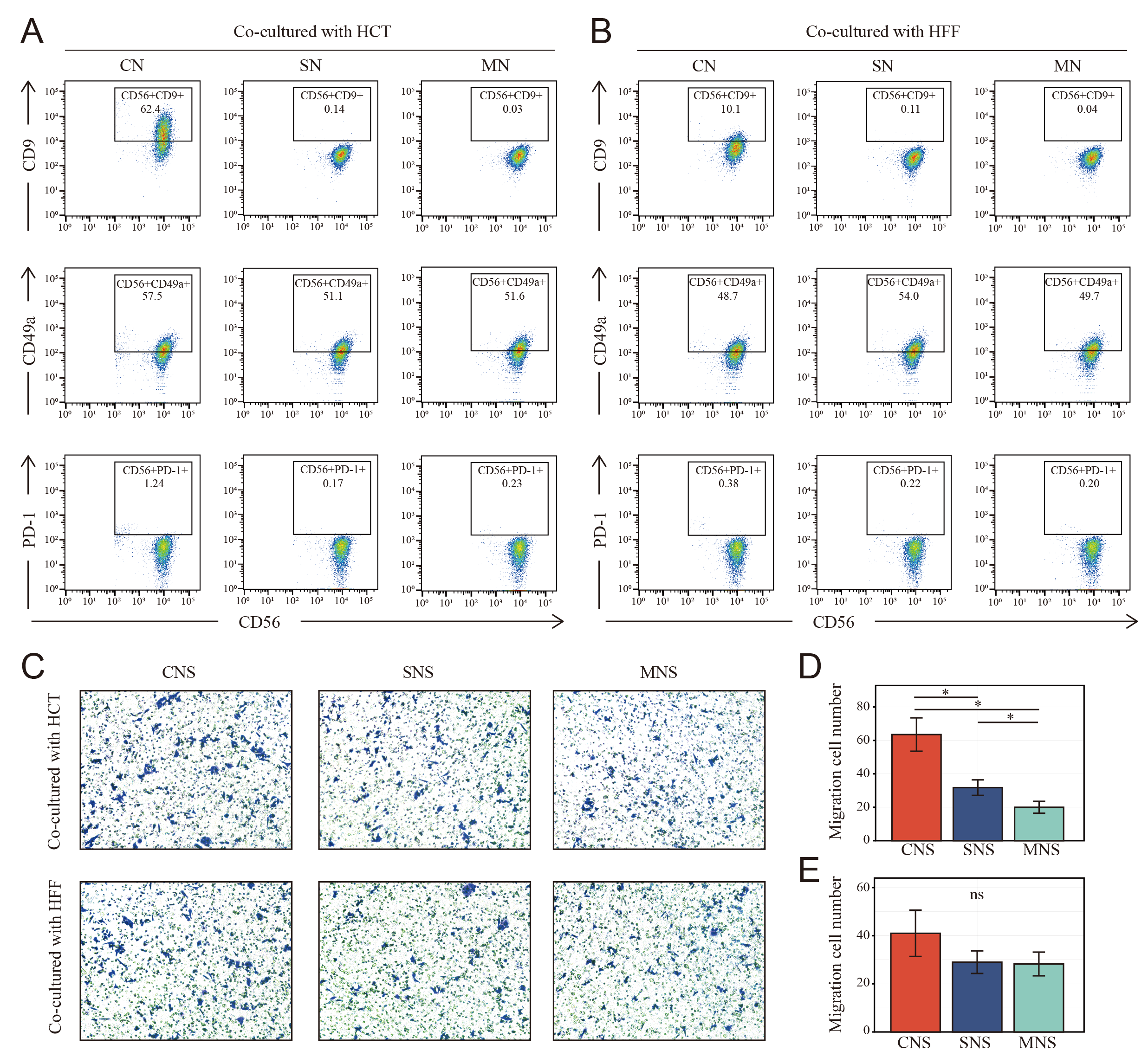

### Figure-S11

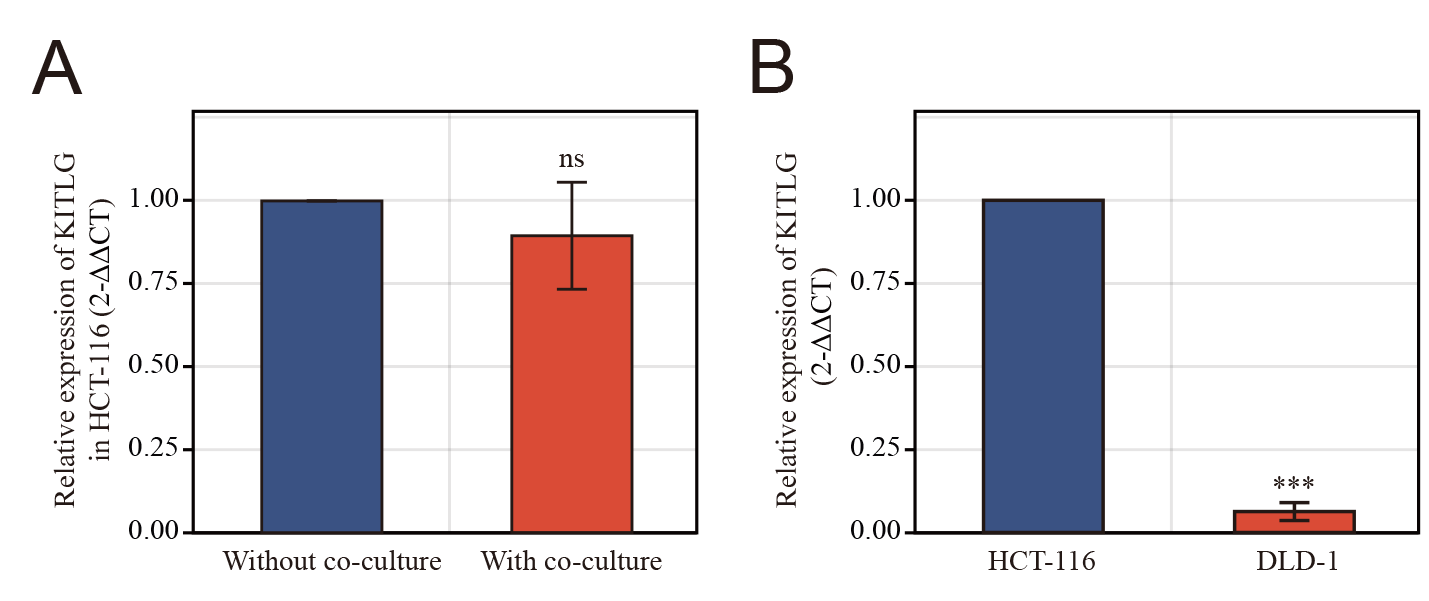
