## Supplement Tables for "Resting natural killer cells promote the progress of colon cancer liver metastasis by elevating tumor-derived sSCF"

Table S1. Fluorescence Minus One control of CD56, CD9, PD-1 and CD49a.

| Group | CD56-APC | CD9-FITC | CD49a-PerCP | PD-1-PE |
| --- | --- | --- | --- | --- |
| CD56-FMO | - | + | + | + |
| CD9-FMO | + | - | + | + |
| CD49a-FMO | + | + | - | + |
| PD-1-FMO | + | + | + | - |

Table S2. Fluorescence Minus One control of KIR2DL4 and GZMK.

| Group | CD56-APC | KIR2DL4-PE | GZMK -FITC |
| --- | --- | --- | --- |
| KIR2DL4-FMO | + | - | + |
| GZMK -FMO | + | + | - |

Table S3. Primers for qPCR.

| Target gene | Sequence |
| --- | --- |
| KITLG Forward primer | CAGAGTCAGTGTCACAAAACCATT |
| KITLG Reverse primer | TTGGCCTTCCTATTACTGCTACTG |
| GAPDH Forward primer | GGGGAGCCAAAAGGGTCATCATCT |
| GAPDH Reverse primer | GACGCCTGCTTCACCACCTTCTTG |

Table S4. Characterized genes for Single-cell transcriptomic analysis

| **T cells** | **NK cells** | **Native B cell** | **Monocyte** | **Plasma cell** | **Neutrophil cell** | **Monocyte CD16-** | **Monocyte CD16+** |
| --- | --- | --- | --- | --- | --- | --- | --- |
| IL7R | NKG7 | CD79A | SPP1 | IGKC | CXCL8 | S100A9 | C1QA |
| LTB | CCL5 | MS4A1 | APOE | JCHAIN | G0S2 | VCAN | MARCO |
| SPOCK2 | KLRD1 | HLA-DRA | APOC1 | IGLC2 | C15orf48 | S100A8 | C1QB |
| KLRB1 | XCL2 | VPREB3 | TIMP1 | IGHA1 | FCGR3B | LYZ | SELENOP |
| TRAC | CMC1 | BANK1 | IL1B | IGHA2 | NAMPT | S100A12 | PDK4 |
| SARAF | GZMB | IGHD | CXCL2 | IGLC3 | S100A8 | FCN1 | CST3 |
| LDHB | GZMH | CD83 | IFI30 | IGHG1 | BCL2A1 | MNDA | CD5L |
| RORA | CST7 | CD74 | C1QC | IGHG3 | S100A9 | THBS1 | SLC40A1 |
| NDFIP1 | GZMA | HLA-DQB1 | CXCL3 | IGHG4 | CSF3R | CYP1B1 | VCAM1 |
| CD2 | IFNG | MEF2C | LYZ | MZB1 | SOD2 | CTSS | C1QC |
| LEPROTL1 | GZMK | HLA-DPB1 | CTSD | SSR4 | MNDA | GCA | MS4A7 |
| CD3D | CCL4 | CD37 | CST3 | DERL3 | SMIM25 | CD14 | MS4A6A |
| TPT1 | CTSW | TNFRSF13C | FTL | TNFRSF17 | SLC25A37 | IL1R2 | CD163 |
| RPS3A | GZMM | HLA-DQA1 | C1QB | SEC11C | PTGS2 | CD163 | LIPA |
| CD3E | CD8B | HLA-DPA1 | CCL3 | XBP1 | IFITM2 | PLBD1 | SDC3 |
| RPS20 | PRF1 | HLA-DRB1 | C1QA | HSP90B1 | CMTM2 | SERPINA1 | CTSB |
| GPR183 | CD8A | CD79B | HLA-DRA | PRDX4 | CCL3L1 | CSTA | FCGRT |
| TRBC2 | HCST | LINC00926 | CTSB | HERPUD1 | ALOX5AP | SLC11A1 | FTL |
| RPL22 | PIK3R1 | RALGPS2 | HLA-DRB1 | TXNDC5 | LST1 | FPR1 | HMOX1 |
| RPL34 | CCL4L2 | LY9 | CCL3L1 | FKBP11 | NEAT1 | SERPINB1 | CFD |

Table S5. Characterized genes for spatial transcriptomic analysis

| **Tumor** | **Normal epithelium** | **Hepatocytes** | **Lamina propria** | **Fibroblast** | **Smooth muscle** |
| --- | --- | --- | --- | --- | --- |
| SCD | SELENOP | APOA1 | JCHAIN | COL1A1 | CSRP1 |
| RPL4 | CKB | APOA2 | DERL3 | COL4A1 | AOC3 |
| LRRC75A | TSPAN1 | ALB | MZB1 | COL3A1 | TPM2 |
| FABP1 | MT-CO1 | HP | IGHA1 | SPARC | MYL9 |
| CSTB | ZG16 | FGB | IGHG4 | COL5A1 | MYH11 |
| GGCX | SLC26A2 | RBP4 | TXNDC5 | COL4A2 | SELENOM |
| VEGFA | KRT19 | FGG | IGHA2 | C3 | CNN1 |
| ID1 | FCGBP | AHSG | IGHG1 | MCAM | TAGLN |
| TNFRSF6B | SDCBP2 | CYP3A4 | IGHG3 | IGFBP7 | FLNA |
| RPL34 | HLA-C | APOC3 | IGKC | CD93 | ACTG2 |
| S100A6 | B2M | AMBP | CD27 | ENG | ACTA2 |
| HMGCS1 | HLA-B | ORM1 | C3AR1 | COL5A2 | GREM2 |
| ZFAS1 | PLAC8 | CYP2E1 | TRAC | FSTL1 | DES |
| TGFBI | ADM | HRG | TYMP | CTSK | CALD1 |
| IFITM1 | MXD1 | GC | LTB | SERPINF1 | FLNC |
| CCDC88B | ITLN1 | A1BG | JAK3 | MMP11 | MYLK |
| ATP1B1 | GUCA2A | FGA | TRBC2 | MMP2 | SYNPO2 |
| RPL22L1 | MALL | ORM2 | TBC1D10C | C1R | ANGPTL2 |
| CD55 | SECTM1 | ST6GAL1 | CORO1A | COL1A2 | TNS1 |
| AC092069.1 | MYO15B | MT-ATP8 | CXCR4 | HSPG2 | PDE5A |

Table S4. Characterized genes of NK subsets

| **Cluster 0** | **Cluster 1** | **Cluster 2** | **Cluster 3** | **Cluster 4** | **Cluster 5** | **Cluster 6** | **Cluster 7** |
| --- | --- | --- | --- | --- | --- | --- | --- |
| GZMK | FGFBP2 | EEF1G | FCER1G | MTRNR2L12 | IGHA1 | MT-ND6 | CXCL13 |
| IL7R | GNLY | EGR1 | CCL3 | PLCG2 | IGKC | PPP1R1B | CTLA4 |
| TUBA4A | FCGR3A | RPL17 | AREG | HLA-DRB5 | LDLRAD4 | RNF19A | RBPJ |
| CD8A | PTGDS | RNASEK | TYROBP | PSMB9 | IGHA2 | EZR | DUSP4 |
| CXCR4 | GZMB | ATP6V0C | CEBPD | XCL2 | IGLC2 | HLA-DQA1 | TNFRSF18 |
| CD8B | SPON2 | KLRK1 | XCL1 | HIST1H4C | IL7R | SLC2A3 | GZMB |
| RGCC | PRF1 | LIME1 | KLRF1 | AC007952.4 | ITGA1 | CREM | PHLDA1 |
| RPS20 | MYOM2 | HSPA1B | KLRB1 | RPS26 | SPRY1 | METRNL | LINC01480 |
| RPS2 | NKG7 | JUN | NFKBIA | HCST | JCHAIN | PTPRC | SRGAP3 |
| RPL13A | CLIC3 | EIF4A1 | CLIC3 | CD160 | TMIGD2 | PDE4B | LAYN |
| TRGC2 | EFHD2 | AL627171.2 | GADD45B | MTRNR2L8 | CD55 | LCP1 | FAM3C |
| RPL23A | LAIR2 | NME2 | CMC1 | GIMAP7 | CD52 | NCL | SAMSN1 |
| HLA-DPB1 | GZMH | FOSB | IRF8 | MYL12A | ANKRD28 | YWHAZ | TNFRSF9 |
| ZFP36L2 | PLAC8 | CD69 | IL2RB | ITGB2 | MT-CYB | IGKC | SNX9 |
| HLA-DRB1 | TYROBP | NR4A1 | TXK | Z93241.1 | CAPG | IGHG3 | NR3C1 |
| YBX3 | ADGRG1 | KLRC3 | GSTP1 | TMEM107 | PDE4D | TPM3 | ENTPD1 |
| LYAR | CD247 | MIF | KLRC1 | PSME2 | SMIM3 | REL | CXCR6 |
| RPS16 | CX3CR1 | CRIP1 | CD160 | ATP5F1E | LINC01871 | GRB7 | CREM |
| TRAT1 | KLRF1 | DNAJB1 | GRASP | CORO1A | IGLC3 | YPEL5 | CCL20 |
| CRTAM | HOPX | BCL11B | TLE1 | PLEKHF1 | TNFAIP3 | HIST1H1E | GAPDH |

Table S5. Characterized genes of NK of different status

| **Resting NK cell** | **Activated NK cell** | **Other NK cell** |
| --- | --- | --- |
| MTRNR2L12 | EGR1 | CXCL13 |
| FGFBP2 | IGHA1 | CTLA4 |
| TYROBP | EEF1G | RBPJ |
| FCER1G | IGKC | DUSP4 |
| NKG7 | RNASEK | TNFRSF18 |
| FCGR3A | ATP6V0C | GZMB |
| CLIC3 | RPL17 | PHLDA1 |
| KLRF1 | LDLRAD4 | LINC01480 |
| GZMK | IGHA2 | SRGAP3 |
| PLAC8 | KLRK1 | LAYN |
| SPON2 | FOSB | FAM3C |
| PLCG2 | LIME1 | SAMSN1 |
| RPS26 | ITGA1 | TNFRSF9 |
| CMC1 | MT-ND4L | SNX9 |
| ITGB2 | JUN | NR3C1 |
| CD247 | TNFAIP3 | ENTPD1 |
| CST7 | AL627171.2 | CXCR6 |
| GNLY | EIF4A1 | CREM |
| CCL3 | LINC02446 | CCL20 |
| HSPA6 | MT2A | GAPDH |
